# Second messenger control of mRNA translation by dynamic ribosome modification

**DOI:** 10.1101/2020.02.13.947879

**Authors:** Lucia Grenga, Richard Howard Little, Govind Chandra, Stuart Daniel Woodcock, Gerhard Saalbach, Richard James Morris, Jacob George Malone

## Abstract

Control of mRNA translation is a crucial regulatory mechanism used by bacteria to respond to their environment. In the soil bacterium *Pseudomonas fluorescens*, RimK modifies the C-terminus of ribosomal protein RpsF to influence important aspects of rhizosphere colonisation through proteome remodelling. In this study, we show that RimK activity is itself under complex, multifactorial control by the co-transcribed phosphodiesterase trigger enzyme (RimA) and a polyglutamate-specific protease (RimB). Furthermore, biochemical experimentation and mathematical modelling reveal a role for the nucleotide second messenger cyclic-di-GMP in coordinating these activities. Active ribosome regulation by RimK occurs by two main routes: indirectly, through changes in the abundance of the global translational regulator Hfq and directly, with translation of surface attachment factors, amino acid transporters and key secreted molecules linked specifically to RpsF modification. Our findings show that post-translational ribosomal modification functions as a rapid-response mechanism that tunes global gene translation in response to environmental signals.

## Introduction

The plant rhizosphere is a highly complex environment, comprising an intricate spatial organisation of roots and soil, with thousands of competing and cooperating microorganisms linked by dynamic fluxes of nutrients, toxins and signalling molecules (1, 2). Microbial rhizosphere colonisation is a correspondingly complex, multi-stage process by which soil-associated bacteria spatially explore, exploit and defend the root environment (3). The colonisation process begins with chemotaxis to the rhizosphere along an exudate gradient, followed by surface adhesion, migration across the root surface (4) and biofilm formation (5).

Bacterial motility is facilitated by type IV pili, flagella and the production of biosurfactants. Together, these factors enable coordinated swarming motility and rhizoplane migration (4, 6). Conversely, root surface attachment and biofilm formation involve the targeted deployment of exopolysaccharide molecules (5, 7), proteinaceous adhesins (8), and surface lipopolysaccharides (9). In addition to motility and biofilm formation, successful rhizosphere colonisation requires the deployment of enzymes and natural product molecules that fulfil a variety of roles, such as the manipulation of plant growth and behaviour (10–12), metal ion scavenging (13) and antagonism of plant pathogens and other competitors (14, 15). Metabolic adaptation to the rhizosphere environment is also highly important, with subsets of primary metabolic genes and substrate transporters expressed in different rhizosphere environments (16).

To enable effective rhizosphere colonisation, soil bacteria sense many different environmental inputs and translate them into an integrated phenotypic response. This requires an interconnected network of signal transduction systems functioning at multiple regulatory levels, including gene transcription (17), modulation of translational activity (18) and changes in protein function (19). The cyclic-di-GMP (cdG) signalling network mediates the switch between motile and sessile lifestyles in many bacterial species (20) and is a key regulator of rhizosphere colonisation in multiple plant-associated microbes (21–24). CdG signalling in *Pseudomonas* forms a highly complex, non-linear and pleiotropic network, with multiple connections to other signalling systems and phenotypic outputs that vary profoundly in response to environmental cues (25, 26). The model *P. fluorescens* strain SBW25, for example, contains over 40 cdG-metabolic enzymes (27) that influence phenotypes at every regulatory level and whose expression varies throughout rhizosphere colonisation (22). *Pseudomonas* cdG signalling shows extensive overlap with other global gene regulators, such as Gac/Rsm (28, 29) and the RNA-chaperone Hfq (30).

The small hexameric protein Hfq facilitates binding between mRNA and regulatory sRNAs (31, 32) and controls biofilm formation (33), carbon catabolite repression (34), amino-acid transport (35, 36), virulence (37) and motility (36). In *P. fluorescens*, Hfq is important for niche adaptation, with deletion mutants displaying strongly reduced motility, increased surface attachment, and severely compromised rhizosphere colonisation (25, 30). The regulatory connections between cdG and Hfq are reflected in the close phenotypic parallels between mutants in both pathways (30, 38, 39).

We recently identified a further contributor to the post-transcriptional regulatory network in *Pseudomonas* spp. (30). Similar to Hfq and cdG, the ribosomal modification protein RimK controls the transition between active and sessile bacterial lifestyles, with an SBW25 *rimK* mutant affected in motility, root attachment and amino-acid uptake, and showing reduced rhizosphere colonisation efficiency. Deletion of *rimK* in pathogenic *Pseudomonas* spp. leads to significantly reduced virulence and cytotoxicity (30). RimK is an ATP-dependent glutamyl ligase that adds glutamate residues to the C-terminus of ribosomal protein RpsF, which in-turn induces specific changes in the bacterial proteome (30). RimK activity is itself controlled by binding to cdG and the small proteins RimA and RimB. RimK exerts at least some of its regulatory activity indirectly through Hfq, with reduced Hfq levels observed in an SBW25 Δ*rimK* background. Again, the regulatory connection in *Pseudomonas* between RimK, Hfq and cdG signalling leads to extensive overlap between their associated phenotypes (25).

Our research to date leads to a model connecting RimK glutamation of RpsF with proteomic changes that enable environmental adaptation by *Pseudomonas*. Nonetheless, at this stage several key questions remain unanswered. The Rim pathway clearly responds to environmental cues that vary during rhizosphere colonisation (30). However, the nature of these signals and mechanisms by which they control RimK activity are currently unknown, and the relationship between the three Rim proteins and cdG is poorly defined. How RimK modification of RpsF induces downstream effects in the *Pseudomonas* proteome is also not well understood. In particular, we do not understand the extent to which RimK activity is mediated through altered Hfq levels as opposed to other mechanisms, and whether changes in RpsF modification lead directly to altered mRNA translation.

To address these outstanding questions, we used a combination of protein biochemistry, computational modelling, quantitative proteomics and ribosomal profiling (Ribo-seq) to interrogate Rim system function and the relationship between ribosomal modification and changes in the SBW25 proteome. We show that RimK controls proteome remodelling by two distinct routes. Indirectly, the Rim system induces media-dependent changes in Hfq abundance, as described previously (30). Independent of these Hfq-mediated effects, RpsF glutamation leads directly to altered translation of a subset of genes including surface attachment factors and amino-acid metabolism. Rim-induced proteomic changes occur rapidly when bacteria are exposed to Rim activating conditions, confirming that ribosomal modification by RimK functions as a previously uncharacterised, actively regulated mechanism enabling rapid proteome adaptation in response to changes in the rhizosphere environment.

## Results

### Expression of the *rimABK* operon is controlled by temperature and nutrient availability

Our previous analysis showed that expression of the *P. fluorescens* SBW25 *rim* operon changes throughout rhizosphere colonisation, peaking in the initial stages of colonisation and then dropping dramatically as the root microbial population becomes established (30). This suggested that *rim* expression was subject to control by uncharacterised signals from the rhizosphere environment and prompted us to investigate the *rimABK* operon in more detail. First, RT-PCR was used to show that the three *rim* genes are transcribed as a single polycistronic operon (Fig S1A), whose transcriptional start site was mapped to 28-30 bp upstream of the *rimA* ORF using 5’ RACE (Fig S1B). To examine the relationship between *rim* expression and environmental signals in more detail, we used qRT-PCR to quantify *rimK* mRNA abundance in SBW25 exposed to a variety of different nutrient conditions and abiotic stresses. Significant increases in *rimK* expression were observed for cells transferred to low temperature (8°C), and exposed to nutrient-limiting conditions (Fig 1), consistent with earlier findings that *rim* expression is reduced in the established (nutrient-replete) wheat rhizosphere (30).

**Figure 1:**
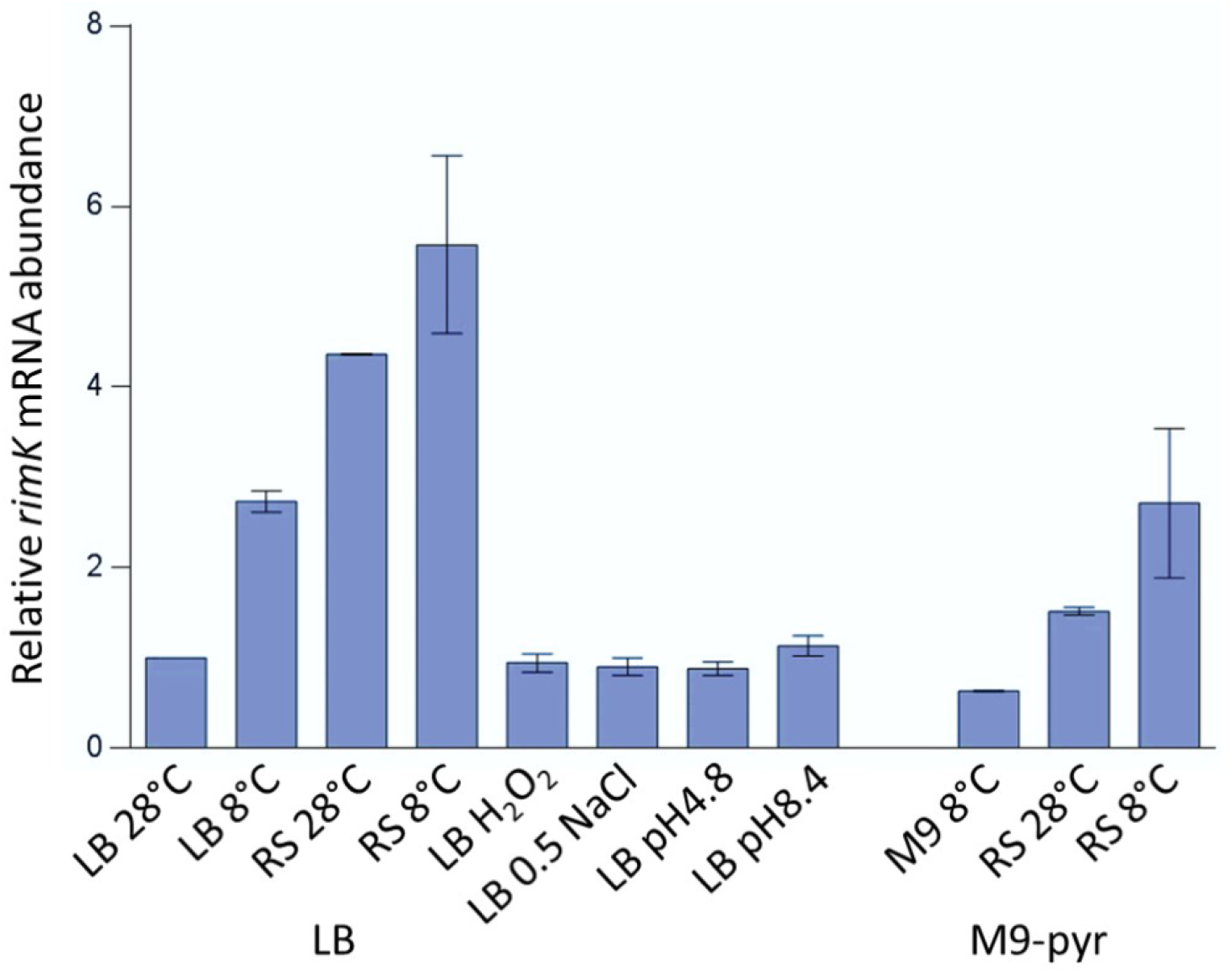
*rimK* expression is stimulated by low temperature and nutrient starvation. mRNA abundance (qRT-PCR data) relative to cells grown overnight in LB and transferred to LB at 28 °C for 45 minutes. In each case cells were grown in LB or M9-pyruvate and transferred to a new set of conditions for 45 minutes prior to sampling. RS – carbon free ‘Rooting Solution’ (30), H_2_O_2_ – LB containing 1.0 mM H_2_O_2_, LB 0.5 NaCl – LB containing 0.5 M NaCl. Data are presented +/− the standard error of three replicates. The experiment was repeated three times independently and a representative is shown.

### RimA and cdG stimulate RimK ATPase activity and RpsF glutamation

Our initial study suggested that a major element of RimK regulation is mediated post-transcriptionally, by interaction with RimA, RimB and the second messenger molecule cdG (30). All three regulators were shown to bind directly to RimK, whose ATPase activity increases upon addition of either protein or cdG *in vitro*. However, effects on RpsF glutamation were only examined (and shown to increase) for addition of cdG (30). At this stage, how the RimABK-cdG system functions to control ribosomal protein modification remains unknown. To address this, we developed kinetic models of the system and conducted a series of biochemical experiments, measuring the effects of each regulator individually and in combination on RimK activity.

The effects of both RimA and cdG on RimK ATPase activity were dependent on a roughly equimolar ratio of protein/messenger with RimK (Fig 2A and 2E). Combined with earlier observations (30), this strongly suggests that RimK activity is controlled by direct interaction in each case. A simple explanation for the change in RimK activity is that RimK can exist in two conformational states, one with basal activity and one with higher activity. To evaluate this hypothesis, we built models of protein binding based on mass action kinetics for the individual reactions (Table S1). The two-state model captures the behaviour of each of the individual experiments, however, the predicted differences in the equilibrium binding constants are not consistent with existing data (Fig S2). Using K_d_ = 1 µM for cdG binding to RimK (30) leads to 96% of RimK being in a cdG bound state under the experimental conditions and predicts a marginal ATPase activity increase upon addition of RimA for a two state model. Experimental validation qualitatively confirms this increase but quantitatively falsifies the two-state model, showing that the effects of RimA and cdG are approximately additive (Fig 2E). This suggests that RimK can exist in at least four different states with varying ATPase activities (Table S1). This four-state model of RimK ATPase activity provides a good quantitative fit with the experimental data as well as reasonable K_d_ values (Fig S2).

**Figure 2:**
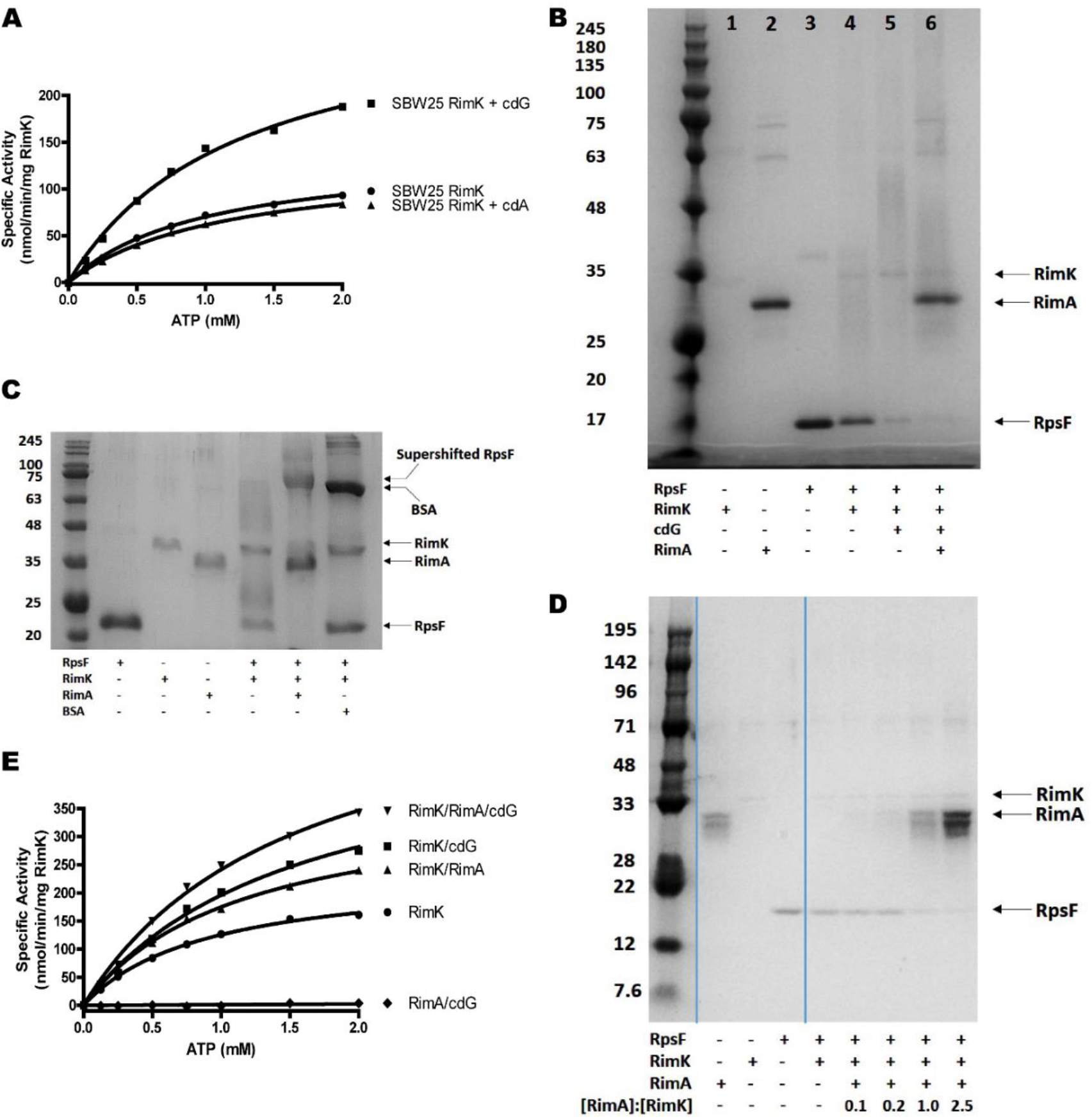
(A) CdG stimulates RimK ATPase activity. RimK was present at a concentration of 2.5 µM. Nucleotides were present at a concentration of 25 µM. **(B) CdG and RimA additively accelerate the rate of RpsF modification.** 12.5% SDS-PAGE gel. *E. coli* RpsF, SBW25 RimK and SBW25 RimA were present at a concentration of 6.4 µM, 3.8 µM and 3.9 µM respectively. The positions of RimK, RimA and RpsF are indicated. CdG was present at a concentration of 150 µM. The samples were incubated overnight prior to electrophoresis. **(C) RimA alone accelerates the rate of RpsF modification.** 12.5% SDS-PAGE gel. *E. coli* RpsF was present at a concentration of 13.6 µM and SBW25 RimK and RimA were present at a concentration of 3.8 µM and 3.9 µM respectively. The position of a super-shifted RpsF band in the penultimate lane and the positions of RimK, RimA, RpsF and BSA are also indicated. The samples were incubated overnight prior to electrophoresis. **(D) Equivalent concentrations of RimA are required to stimulate RimK modification activity.** Samples were run on a 12.5% SDS-PAGE gel. *E. coli* RpsF was present at a concentration of 6.8 µM, SBW25 RimK at 1.9 µM. RimA was present at the indicated ratio relative to RimK in each case. The positions of RimK, RimA and RpsF are indicated. Samples were incubated for 60 minutes prior to electrophoresis. **(E) The ATPase activity of RimK is stimulated additively by RimA and cdG.** SBW25 proteins were present at 1 µM. CdG was present at a concentration of 25 µM.

To determine whether RimK ATPase activity was a good proxy for RimK glutamate ligase activity on RpsF we extended our experiments to include RpsF. In agreement with our earlier work (30), cdG was shown to stimulate both ATPase activity and RpsF modification (Fig 2A, B). This stimulation was highly specific, as incubation with the dinucleotide signalling molecule cyclic-di-AMP had no effect (Fig 2A). RimA addition also directly and specifically boosted RimK enzyme activity (Fig 2C, D). Furthermore, this stimulation was additive with that provided by cdG (Fig 2B, Lane 6).

### RimB stimulates RimK ATPase activity, but suppresses RpsF glutamation

To investigate the effect of RimB, we incubated RimK with RimB and observed a marked increase in RimK ATPase activity, beyond that achieved by addition of RimA and cdG (Fig 3A, (30)). Based on our experiments with RimA and cdG, we predicted that increased ATPase activity would translate to enhanced glutamate ligase activity. Interestingly however, this was not the case. Not only was the observed ATPase activity increase largely abolished by addition of RpsF to the reaction (Fig 3A), RimB addition produced a strong suppressive effect on the ability of RimK to shift the RpsF band in our glutamation assays (Fig 3B, (40)). A possible explanation is that RimB blocks access of RimK to RpsF. To test this hypothesis we measured the effect of different concentrations of RimB. The prediction is that comparable amounts of RimB would be required to largely saturate RimK and render it inactive for RpsF glutamation. We found that the suppressive effect was indeed strongly dependent on the concentration of RimB in the reaction (Fig 3C) but that surprisingly small concentrations of RimB were sufficient to reduce glutamation, arguing against a stoichiometric binding model of RimK inhibition. This decrease could only be partially countered by increasing glutamate concentration (Fig 3C), eliminating glutamate availability as the cause of RimB suppression of RpsF band-shifting under our test conditions. Thus, RimA, RimB and cdG enhance RimK ATPase activity, and RimA and cdG enhance but RimB suppresses RimK glutamylation activity.

**Figure 3:**
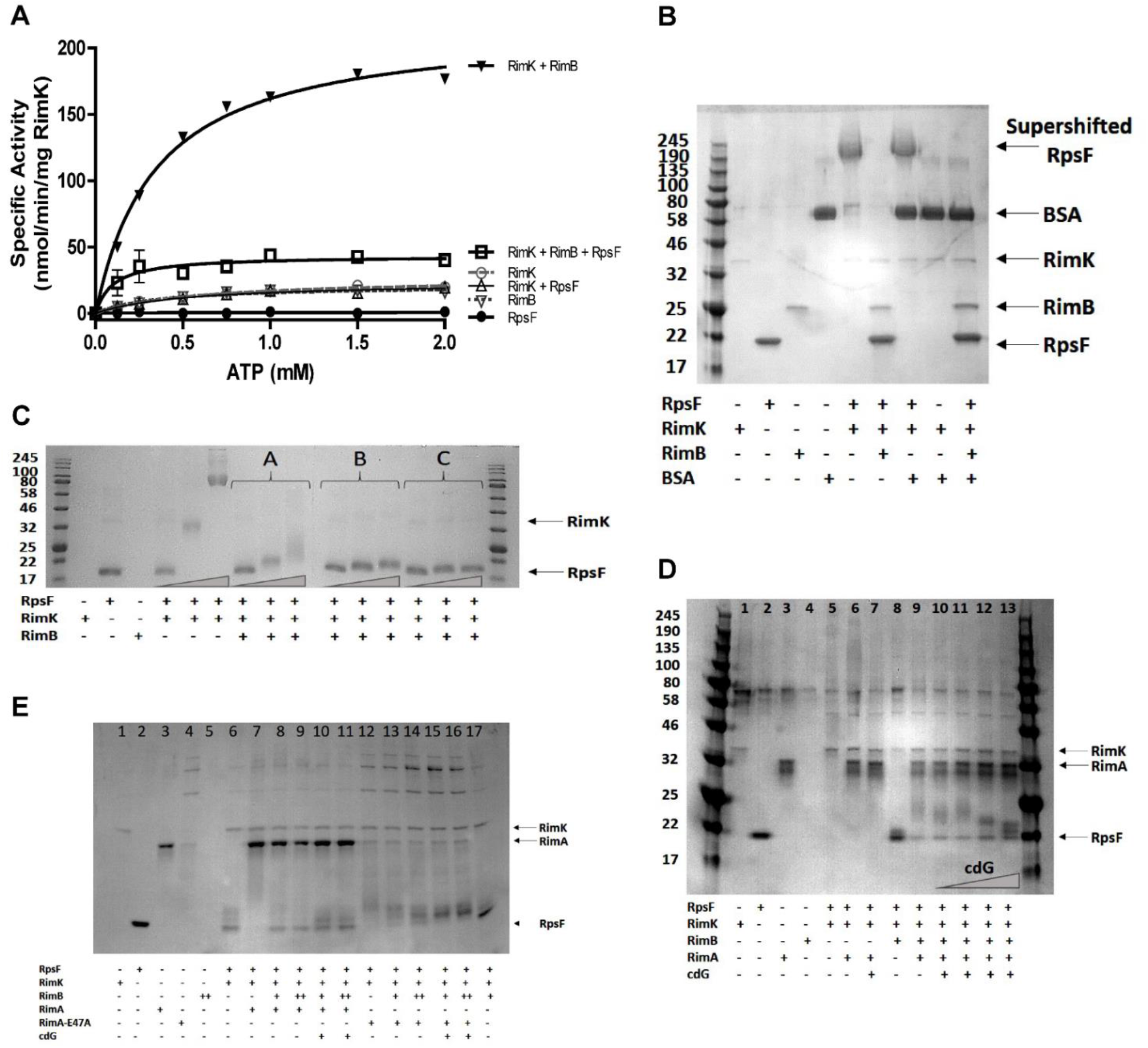
(A) The presence of RpsF drastically reduces the stimulation of RimK ATPase activity by RimB. *E. coli* RpsF, SBW25 RimK and SBW25 RimB were present at a concentration of 1 µM. **(B) RimB specifically antagonizes RimK modification activity.** 12.5% SDS-PAGE gel. *E. coli* RpsF was present at a concentration of 6.8 µM, BSA was present at a concentration of 3.8 µM, SBW25 RimK and RimB were present at a concentration of 7.5 µM and 12 µM respectively. The positions of the supershifted RpsF band, RimK, RpsF, RimB and BSA are indicated. The samples were incubated overnight prior to electrophoresis. **(C) Trace concentrations of RimB are sufficient to significantly antagonize modification of RpsF.** 12.5% SDS-PAGE gel. *E. coli* RpsF and SBW25 RimK were present at a concentration of 6.8 µM and 3.8 µM respectively. The ratio of RimB to RimK in lanes labelled A, B and C was 1:3, 1:16 and 1:62.5 respectively. The grey triangles represent increasing glutamate concentrations of 0.2 mM, 2 mM and 20 mM. The positions of RimK (barely visible at this concentration) and RpsF prior to modification are indicated. **(D) CdG attenuates the ability of RimA to stimulate RimK modification activity in a RimB-dependent manner.** 12.5% SDS-PAGE gel. *E. coli* RpsF, SBW25 RimK, SBW25 RimA and SBW25 RimB were present at a concentration of 6.8 µM, 3.8 µM, 3.9 µM and 6.0 µM respectively. The blue triangle represents increasing cdG concentrations of 5, 10, 50 and 200 µM. Lanes 9 and 11 (containing WT RimA) show increasing densities of low molecular weight RpsF species upon addition of cdG. The positions of RimK, RimA and RpsF (prior to modification) are indicated. **(E) A CdG binding mutant of RimA is able to fully stimulate RimK modification activity in the presence of the nucleotide.** 12.5% SDS-PAGE gel. *E. coli* RpsF, SBW25 RimK and SBW25 RimA proteins were present at a concentration of 6.8 µM, 3.8 µM and 3.9 µM respectively where indicated. RimB was present at either 1.7 µM (denoted by a single cross below the gel) or 3.4 µM (denoted by a double cross below the gel). Cyclic-di-GMP is present at a concentration of 200 µM. Lanes 14 and 16 (containing RimA-E47A) reveal loss of the unmodified RpsF band and increasing intensity of larger species upon cdG addition. The positions of RimK, RimA and RpsF (prior to modification) are indicated.

Interestingly, RimA was able to boost the RpsF modification activity of RimK even in the presence of RimB (Fig 3D, Lanes 8-9). Based on our RimK ATPase data we would expect cdG to boost RpsF modification even further. Puzzlingly, increasing concentrations of cdG actually attenuated the effect of RimA, evidenced by an increasing concentration of smaller molecular weight glutamated RpsF species (Lanes 10-13). This effect appeared to be dependent on the presence of RimB as no attenuation of RimA stimulation of RimK modification activity was observed in the absence of RimB (Lanes 6 and 7). The suppressive effect of cdG on RimK behaviour was dependent on the phosphodiesterase activity of RimA, with an enzymatically inactive RimA variant (RimA-E47A) able to stimulate RimK activity both in the presence or absence of cdG (Fig 3E).

Despite RimB and RimK being co-expressed (Fig S1A) and interacting directly with one another (30), the strong RimB suppression of RimK band-shifting still occurred when the ratio between RimB and RimK was as low as 1:63 (Fig 3C). This strongly argues against a mechanism for RimK regulation based on direct interaction with RimB. This falsifies our initial model in which we suggested that RimB functions as a direct regulator of RimK activity (30) and together with the suppression effect of cdG highlights complexities not accounted for in our stoichiometric kinetic model.

### RimB functions as a specific protease for the modified C-terminal of RpsF

The above observations suggest an enzymatic role for RimB. RimB shows modest secondary structural similarity to Pfam class PF05618 (ATP-dependent zinc proteases), leading us to hypothesize that the apparent suppression of RimK band-shifting may represent cleavage of C-terminal poly-glutamate (poly-E) tails, following their addition by RimK. This would potentially explain how relatively low amounts of RimB are able to effectively prevent RpsF band-shifting by RimK. To test this, we first examined the effect of adding purified RimB to RpsF samples that had been previously glutamated by RimK addition (Fig 4A). Addition of RimB to RimK-modified samples restored the RpsF bands to their original gel positions, consistent with cleavage of the poly-E tail. To exclude the possibility that RimB activity may be mediated through RimK, we purified and incubated with RimB a modified RpsF allele (RpsF_10glu_) with 10 additional C-terminal glutamate residues. The RimB protein was able to reduce the mass of RpsF_10glu_ towards that of WT RpsF, confirming its protease activity (Fig 4B). No effect was seen on RimB incubation with unmodified RpsF, suggesting that RimB acts to remove poly-E tails specifically rather than as a protease for the unmodified RpsF protein.

**Figure 4:**
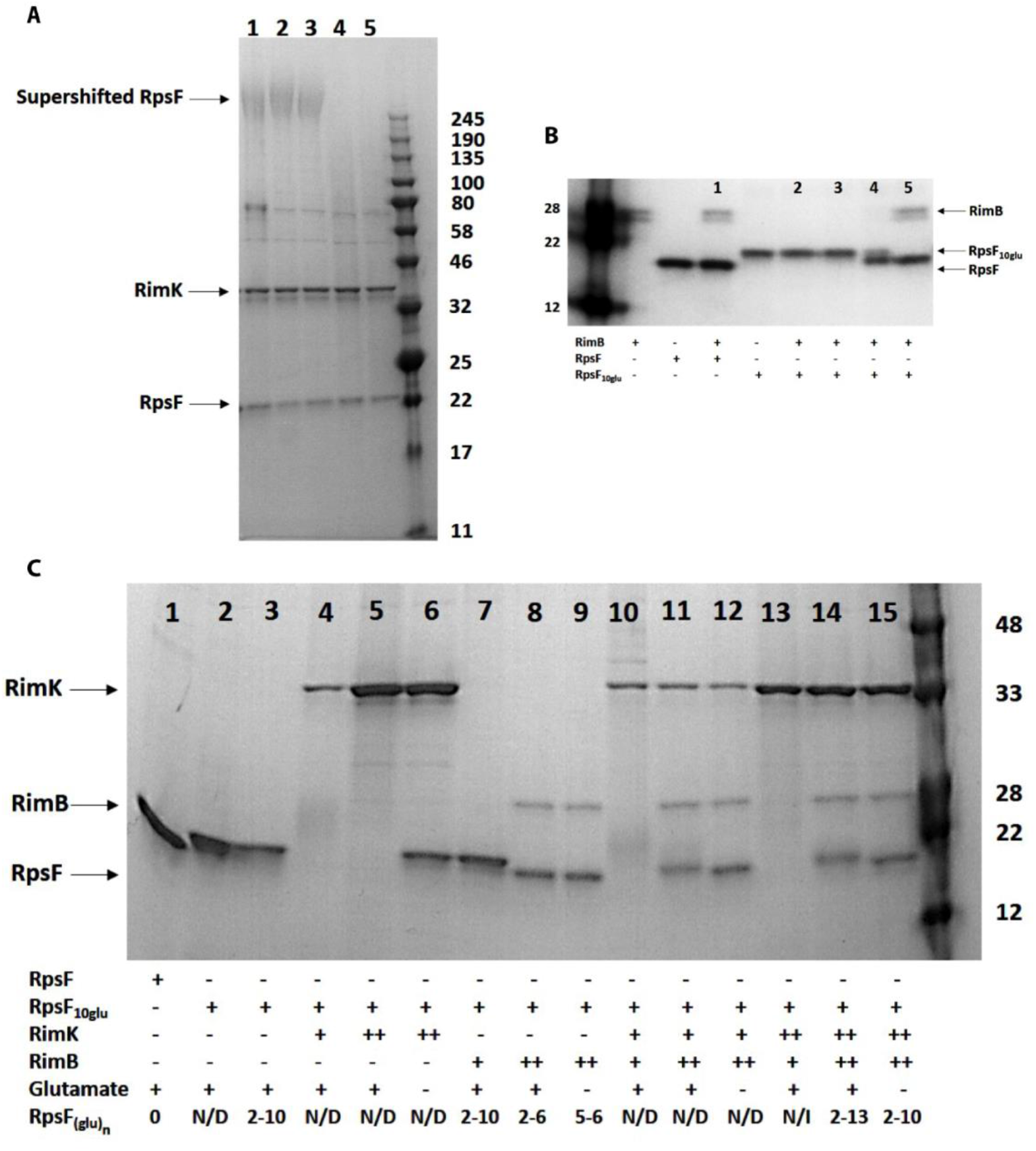
(A) The presence of RimB abolishes the poly-glutamated form of RpsF. 12.5% SDS-PAGE gel. SBW25 RpsF was present at a concentration of 12.8 µM. SBW25 RimK was present at a concentration of 15 µM. Samples were incubated overnight to allow formation of the modified RpsF species shown in Lane 1. The sample was divided into four equal volumes and the following additions made: Lane 1 – buffer only; Lane 2 – 20mM ADP; Lane 3 – 100 nM SBW25 RimB; Lane 4 – 100 nM SBW25 RimB + 20 mM ADP. After 150 minutes incubation, samples were taken for SDS-PAGE analysis. The positions of the supershifted RpsF band, RimK and unmodified RpsF (Lane 1) are indicated. **(B) RimB proteolyzes poly-glutamated RpsF.** 12.5% SDS-PAGE gel. SBW25 RpsF and RpsF_10glu_ were present at a concentration of 12.8 µM. RimB concentrations in the lanes labelled 1-5 were: Lane 1 – 6.0 µM; Lane 2 – 6.0 nM; Lane 3 – 60 nM; Lane 4 – 600 nM; Lane 5 – 6.0 µM. After 5 minutes incubation, samples were withdrawn for SDS-PAGE analysis. The positions of RimB, unmodified RpsF and RpsF_10glu_ are indicated. **(C) RimB cleaves glutamate residues from the C-terminus of RpsF.** 12.5% SDS-PAGE gel. SBW25 RpsF and RpsF_10glu_ were present at a concentration of 12.8 µM. SBW25 RimK was present at a concentration of either 3.8 µM (denoted below the gel as a single cross) or 15 µM (a double cross). SBW25 RimB was present at a concentration of either 60 nM (denoted below the gel as a single cross) or 6.0 µM (a double cross). After 60 minutes incubation, samples were withdrawn for SDS-PAGE analysis. Following SDS-PAGE, gel slices were excised from the region of the gel defined by the unmodified and partially modified RpsF bands present at the base of the gel and submitted for mass spectrometric analysis. N/D = Not Determined; N/I = Nil Identified. The positions of RimK, RimA and RpsF are indicated.

Mass spectrometry analysis of the RpsF bands from the experiment in Fig 4A indicated that RimB protease activity in fact left 4-5 glutamate residues at the RpsF C-terminus. To investigate this activity further, different combinations of RimK, RimA and glutamate were combined *in vitro* and incubated for 60 minutes prior to separation by SDS-PAGE and mass spectrometry analysis to determine the RpsF glutamation state (Fig 4C). As predicted, addition of RimK resulted in unregulated modification of RpsF_10glu_ in a glutamate-dependent manner (Lanes 4-6). Introduction of RimB to RpsF_10glu_ at a concentration of 6 µM resulted in the removal of C-terminal glutamates from RpsF irrespective of the presence of glutamate (Lanes 8 and 9). In the presence of RpsF_10glu_ and RimK, increasing concentrations of RimB partially restored protein density in the RpsF region of the gel (Lanes 10 and 11) Removal of glutamate residues from RpsF by RimB was augmented in the absence of externally added glutamate, presumably due to reduced RimK glutamation activity (Lane 12). In the presence of RpsF_10glu_, 60nM RimB, glutamate and elevated levels of RimK, no RpsF protein was detected in the RpsF region of the gel due to hyper-modification by RimK ((30), Lane 13). Increasing [RimB] to 6 µM restored protein density to the RpsF region of the gel, although glutamate chain lengths of up to 13 residues were still obtained (Lane 14). Removal of glutamate under these conditions reduced the extent of glutamation, but not to the extent of Lanes 8 and 9, indicating that RimK is able to utilise glutamate liberated from RpsF_10glu_ by RimB (Lane 15).

### Computational models postulate that addition and proteolytic removal of glutamates can lead to systemic addition of ∼4 glutamates to many ribosomes

To explain the complex interplay observed in this system, we incorporated protease activity of RimB into a model for RpsF glutamation. We built a stochastic model for chain extension, assuming that without RimB, chain extension is processive and RimK can either add a glutamate or release RpsF. Depending on RimK activity, this can lead to very long glutamate chains. Within the model we set a chain length limit of 100 glutamate units (Fig 5). We then included the action of RimB, cutting the chains from the end, one unit of glutamate at a time, down to the experimentally determined resultant minimal length of four. Unsurprisingly, this results in seemingly futile cycles of RpsF glutamation by RimK followed by cleavage by RimB. The balance between RimK glutamate ligase activity and RimB protease activity determines the dynamics of RpsF glutamate chain lengths and average chain length distribution (Fig 5).

**Figure 5:**
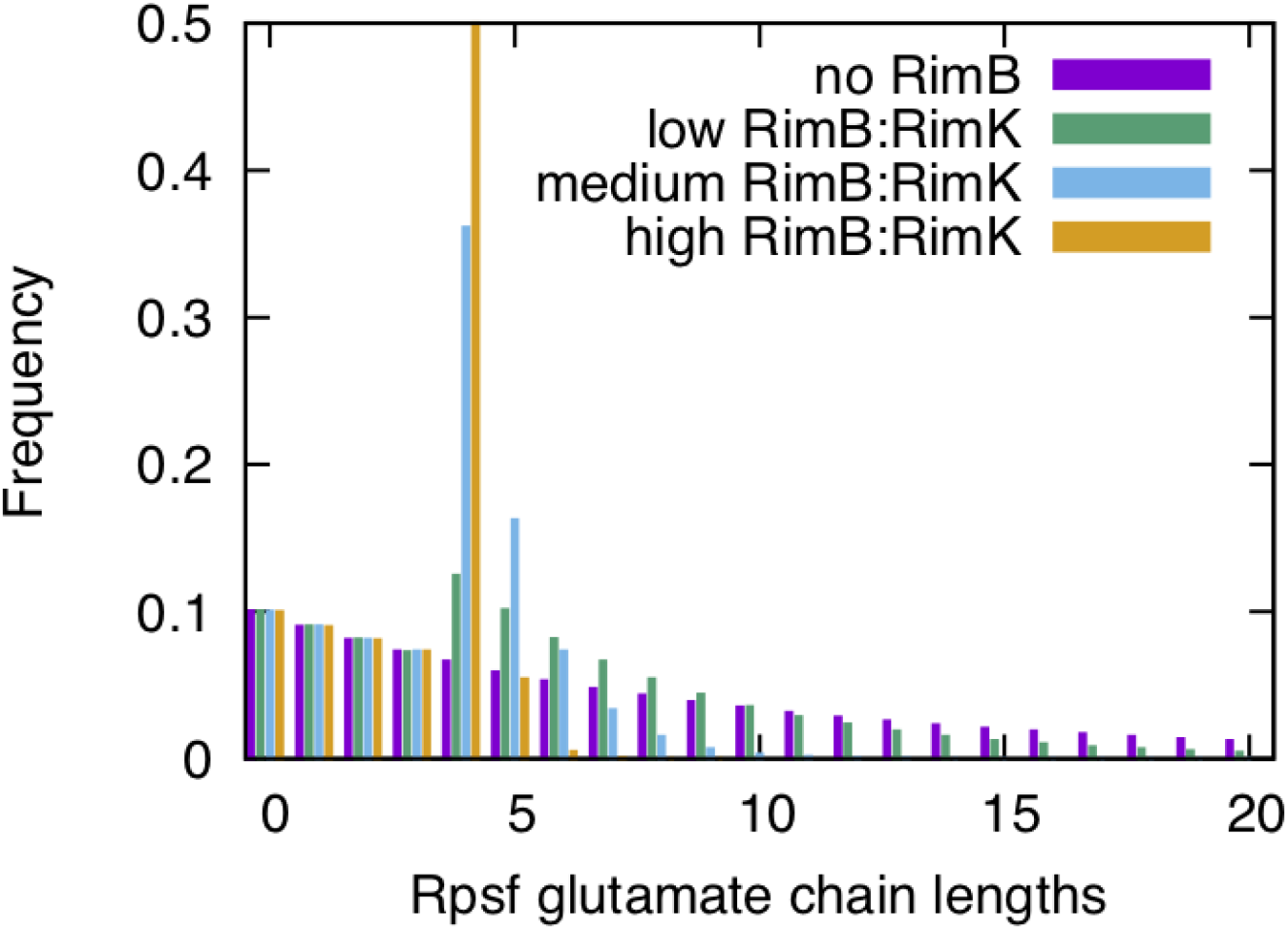
Changing the ratio of protease activity of RimB to RimK glutamate ligase activity changes the glutamate chain length distribution of RpsF. Stochastic simulations of glutamate addition, chain termination and chain cleavage were average over 1,000,000 simulations to obtain the expected chain length distributions as a function of a changing RimB:RimK activity ratio. As has been previously shown, with no dependency of the chain extension and termination probabilities, the chains follow an exponential distribution (purple curve, no RimB protease activity). Introducing RimB protease activity results in longer chains being reduced in length and shifting the distribution towards to minimal length at which RimB is active (four units). Increasing RimB:RimK activity further pushes the chain length distribution to peak sharply around a length of four. By changing the activity of RimK, RimA and cdG can change the ratio of protease to glutamate ligase activity in the system, with an increasing ratio sharpening the chain length distribution around four.

To more realistically model the *in vivo* relationship between the Rim system and the highly abundant RpsF protein, we estimated that a typical cell contains approximately 1 RimABK complex to every 500 RpsF units. This means that in order to understand the overall impact of RimB on the system, we need to consider how the population of RpsF behaves as a function of cdG (i.e. variable RimK activity, assuming RimB activity is constant). Fig 6 shows the distribution of glutamate chain lengths for a population of RpsF molecules. In the absence of RimB, the chain length distribution is broad and importantly we can expect a large number of RpsF proteins to have no glutamate chains at all. With active RimB present, a limited glutamate pool can be effectively equally distributed over all RpsF units in the cell, thus enabling a uniform switch of RpsF. These simulations show the striking results of a changing RimB protease activity vs RimK glutamate ligase activity (governed by cdG) on the number of RpsF units that can be modified. If this global coverage is important, this points to chain lengths of 4 being sufficient to give rise to a change in translation that would allow for a rapid and coordinated response to environmental cues.

**Figure 6:**
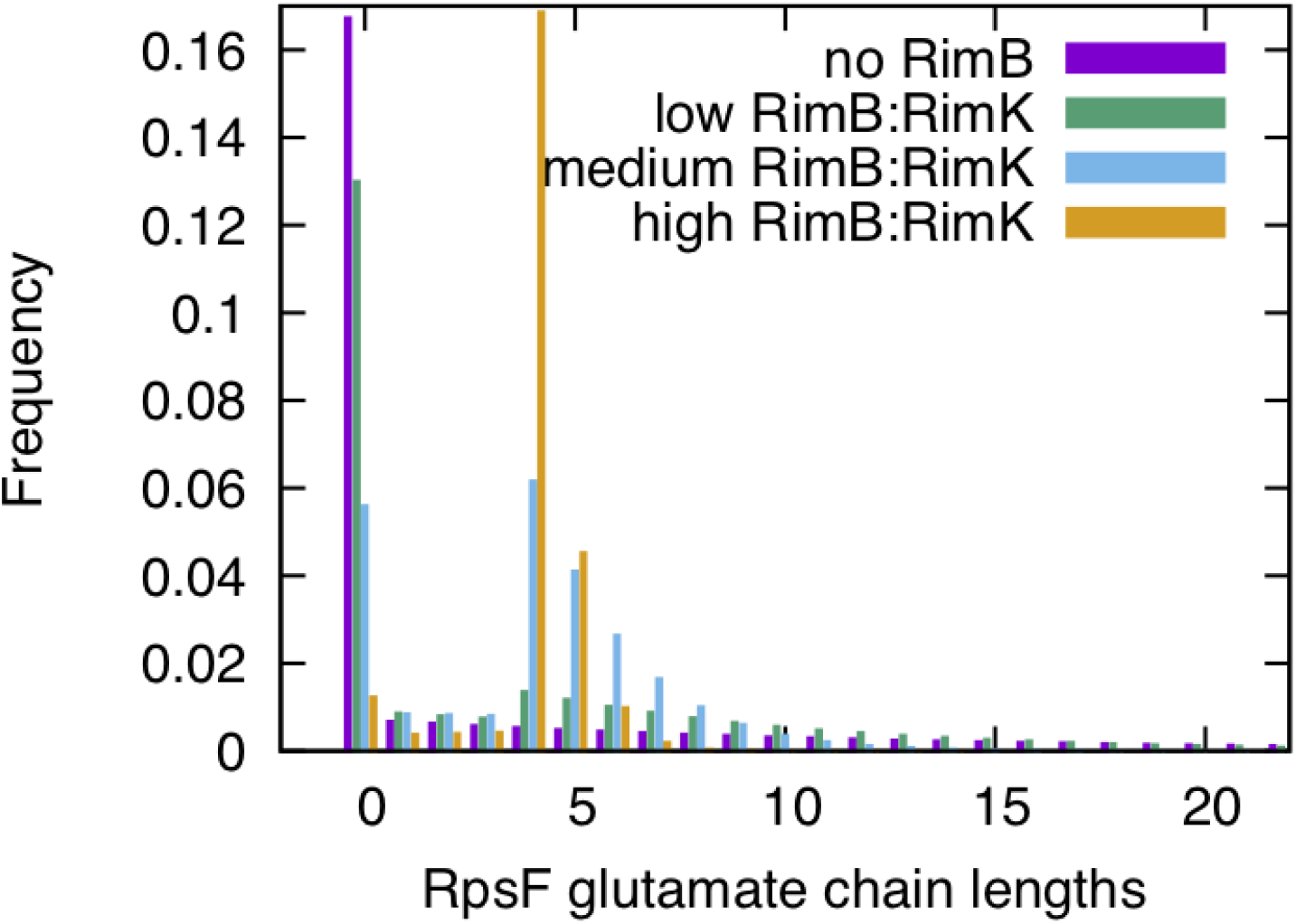
High levels of cyclic di-GMP increase the ratio of RimB protease activity to RimK glutamate ligase activity, leading to a shift towards shorter glutamate chains on RpsF and wider coverage of RpsF modification. For a limited amount of glutamate in the system and the high RpsF to RimABK ratio, the average chain lengths (Fig 5) can lead to drastic changes in the overall modification of ribosomal units. With reduced RimB protease activity vs RimK glutamate ligase activity, a few long chains are produced on some RpsF but the majority have no chains at all (purple, no RimB protease activity). With a higher ratio of RimB protease activity vs RimK glutamate ligase activity, RimK delivers high coverage of shorter chains to more RpsF units (green to blue to orange). This protease to glutamate ligase activity is regulated by cdG and [glutamate]. Stochastic simulations were averaged from 1,000,000 runs using 500 RpsF to every RimABK.

Our *in vitro* data (Fig 4) suggests that RimB proteolytic activity alone cannot restore RpsF to the unmodified state. Thus, to satisfy our model for dynamic ribosomal modification by RimK it is necessary to invoke the presence of a second, ‘reset’ protease that removes post-translational glutamates from RpsF. To seek evidence for additional proteolysis of RpsF, we constructed SBW25 strains in which the *rpsF* gene was extended by 10 glutamate residues. Samples were then grown to exponential or stationary phase and probed with antibodies against RpsF and poly-glutamate (Fig 7). Cells expressing RpsF_10glu_ from an otherwise WT background revealed significant degradation of the RpsF C-terminal tail in comparison to RpsF protein levels under stationary phase conditions where ribosome neogenesis will be severely limited. The loss of the C-terminal poly-glutamate tail was even more pronounced in a Δ*rimK* background. Crucially, cells lacking *rimB* also showed strong reduction of RpsF glutamate tails relative to RpsF levels under stationary phase conditions, consistent with a second, RimB-independent mechanism for RpsF C-terminal degradation.

**Figure 7:**
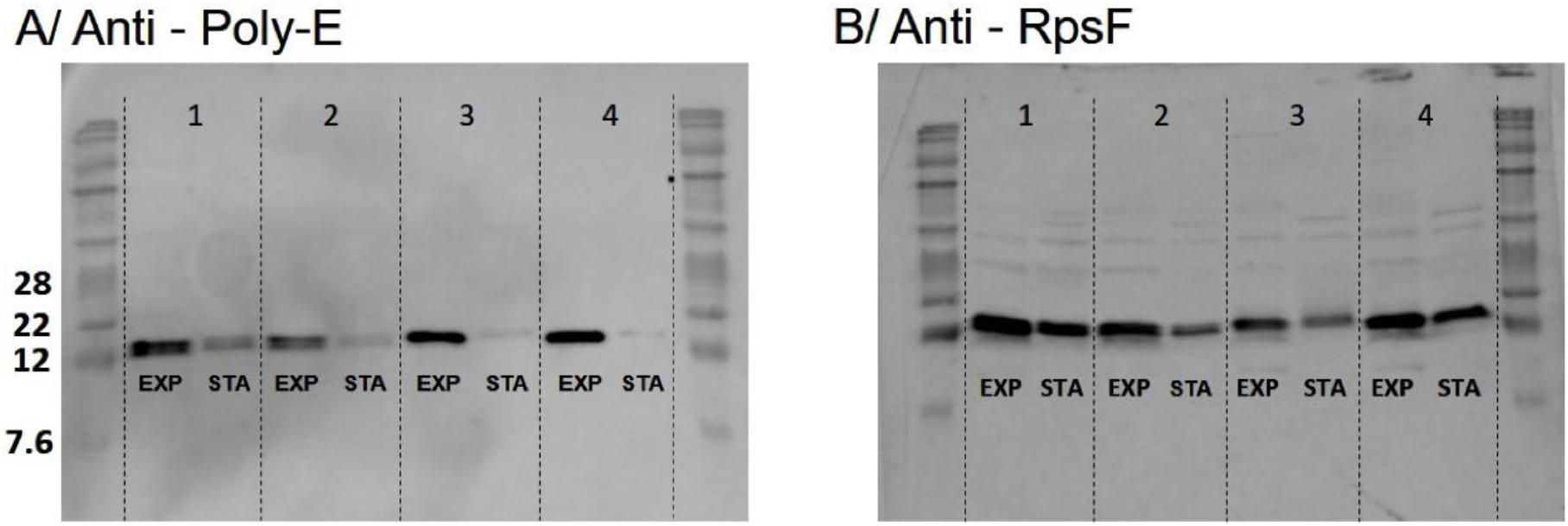
Proteolysis of RpsF glutamate tails is enhanced in stationary phase in a RimB-independent manner. **(A)** Western blot showing levels of RpsF_10glu_ in different genetic backgrounds. Panel 1, SBW25 *rpsF_10glu_*; Panel 2, SBW25 Δ*rimK rpsF_10glu_*; Panel 3, SBW25 Δ*rimB rpsF_10glu_*; Panel 4, SBW25 Δ*rimBK rpsF_10glu_*. **(B)** Western blot showing levels of RpsF protein. The genetic background of panels 1 – 4 is the same as given in (A). EXP denotes samples taken from exponentially growing cells; STA denotes samples taken from stationary phase cells.

### Environmental inputs rapidly affect protein abundance in a RimK dependent manner

To test whether RimK modification of RpsF represents a rapid, active cellular response to environmental signals, we examined the short-term impact of RimK stimulation on the bacterial proteome. Our qRT-PCR data (Fig 1) indicates that *rim* mRNA abundance is promoted in cold, starved cells. Therefore, overnight cultures of WT SBW25 and Δ*rimK* cells grown at 28 °C in LB media were abruptly transferred to 8 °C in nutrient-limiting rooting solution (A), and 28 °C in LB media (B) for 45 minutes. We then conducted a quantitative proteomic analysis experiment using isobaric labelling (iTRAQ) to examine how these conditions affected protein abundance in the WT and mutant backgrounds. When a 2D scatter plot was constructed for this data (Fig 8), proteins whose conditional abundance changes are unrelated to *rimK* deletion sit along a line with gradient +1 (x=y). Deviation from the x=y line indicates an effect of *rimK* deletion on protein abundance under the activating conditions (Fig 8, Table S2), with a substantial fraction of the SBW25 proteome shifting towards higher ratios in Δ*rimK* (Fig 8, and boxplot in Fig S4). Proteins with ratios differing significantly (p≤ 0.05) by a factor ≥2 between WT and Δ*rimK* (located above or below the dashed lines) are coloured in red (up) and blue (down). Substantial agreement with the previously determined RimK regulon (30) could be observed (indicated by the green dots in Fig 8 representing significant differences) confirming that RimK exerts rapid control of the SBW25 proteome under nutrient-limiting conditions and low temperature.

**Figure 8:**
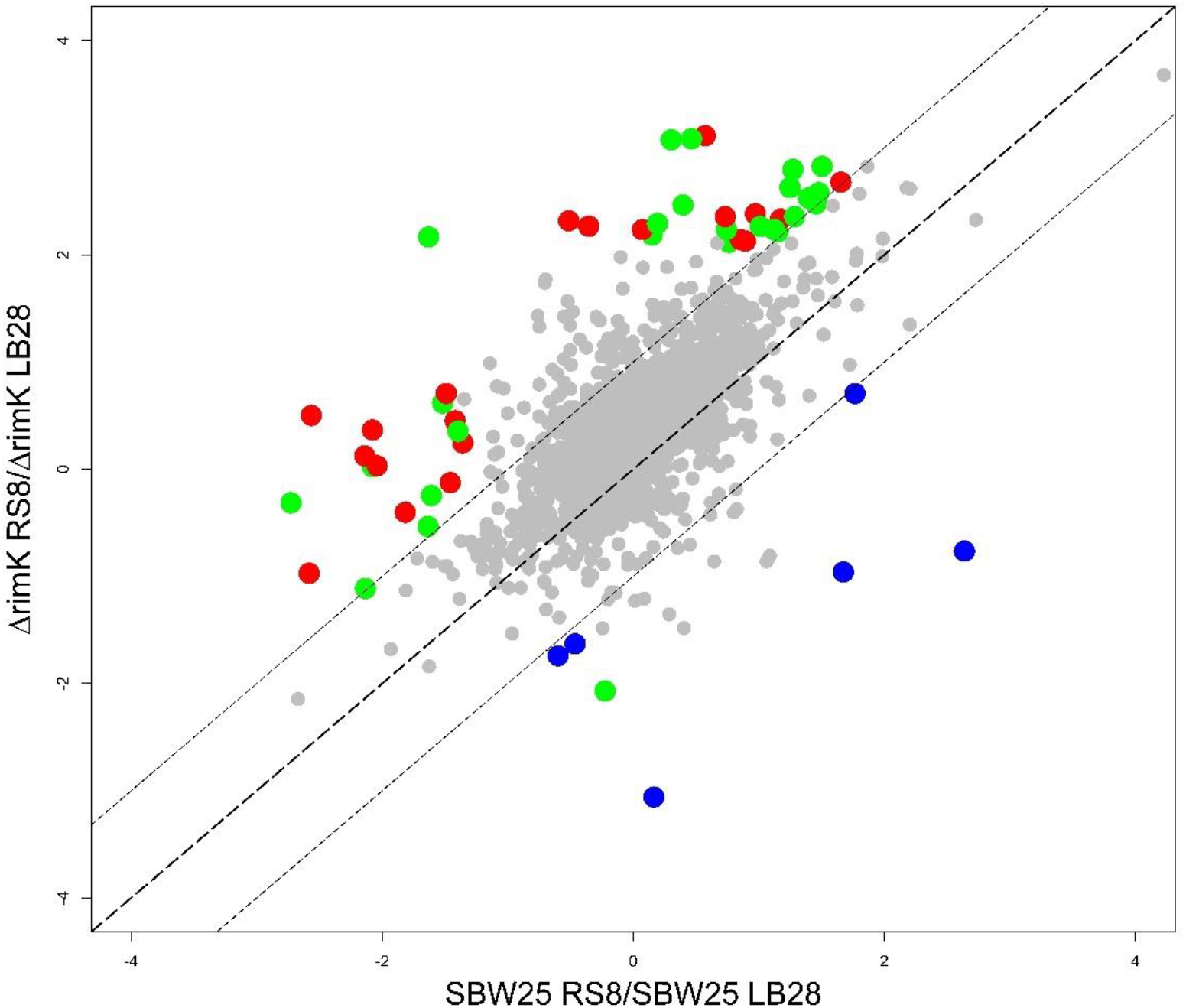
RimK activity rapidly affects the proteome of *P. fluorescens*. 2D scatter plot measuring log_2_-fold change for condition A/condition B for WT on the X-axis and Δ*rimK* on the Y-axis. LB28 denotes 45 minutes incubation in LB medium at 28 °C and RS8 denotes 45 minutes incubation in carbon-free rooting solution at 8 °C. Proteins (filtered for at least unique 2 peptides) whose abundance is significantly (at least 2-fold difference of regulation between WT and Δ*rimK*, at least one p-value ≤ 0.05) affected by *rimK* deletion are highlighted in red (upregulated) and blue (downregulated in Δ*rimK*). Proteins differentially regulated and belonging to the previously determined RimK regulon (30) are highlighted in green.

The timeframe for the proteomic changes observed in our iTRAQ experiment was much too rapid to be a result of the production of new ribosomes (41), strongly suggesting that the proteomic changes we observe are an active response to modification of ribosomally-associated RpsF proteins.

### RimABK exerts direct and indirect effects on mRNA translation through RpsF glutamation

Our data suggests that a large proportion of the observed *rimK* regulon is controlled through RimK-induced shifts in the abundance of other translational regulators, particularly Hfq (30). To understand the contribution made by RimK glutamation to ribosome function, we must first understand the relationship between Hfq abundance and RimK. To assess the contribution of Hfq to the RimK phenotype, we produced *rimK* mutant strains in which the chromosomal *hfq* gene was flag-tagged at the C-terminus. We then quantified Hfq abundance using a Wes Simple Western system (Bio-Techne) in different media conditions. As expected based on previous results (30), Hfq abundance decreased markedly in the Δ*rimK* background for cells grown in M9-pyruvate media. This reduction in Hfq levels was observed for cells grown to early exponential or to stationary phase. Interestingly, the relationship between RimK and Hfq was reversed for cells grown in M9-pyruvate supplemented with Cas-aminoacids. Under these conditions, *rimK* deletion led to a noticeable increase in Hfq abundance in early exponential phase (Fig 9). The results obtained for stationary phase growth in M9-pyr-Cas followed the same trend, with one Δ*rimK* sample showing an increase in Hfq and the second showing little change. This may reflect the depletion of one or more amino-acid sources in the second sample.

**Figure 9:**
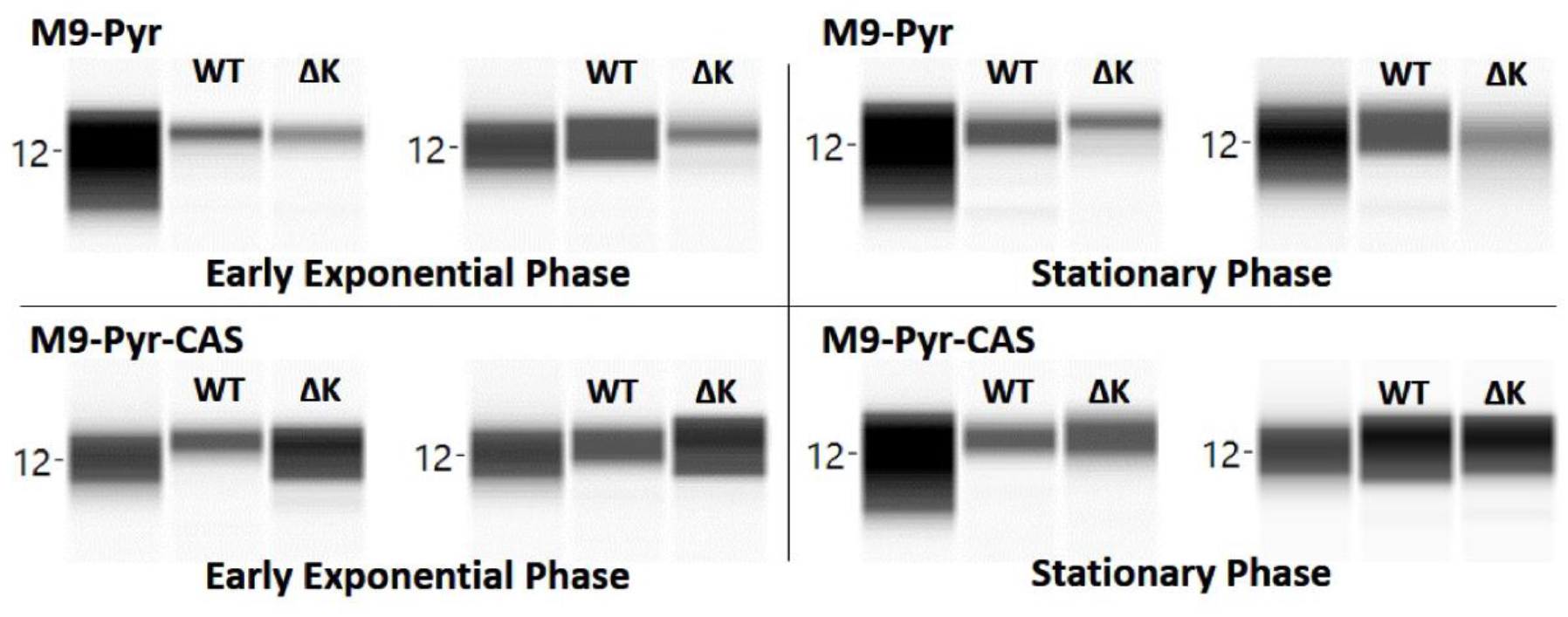
The influence of RimK on Hfq abundance is dependent upon media conditions. SBW25 WT or *ΔrimK* cells were grown in either M9-pyruvate (abbreviated M9-Pyr) or M9-pyruvate with Cas-aminoacids (abbreviated M9-Pyr-CAS) to either early exponential phase or stationary phase as indicated. Each panel shows two biological duplicates; in each case the 12KDa molecular weight marker is shown (Hfq-FLAG has a predicted molecular weight of 12.04 KDa).

Based on these data, we expect that much of the effect of *rim* gene deletion on the SBW25 translatome would be reversed for cells grown with Cas-aminoacids as opposed to with pyruvate alone. To test this, we conducted a Ribo-seq analysis on SBW25 WT and Δ*rimBK* cells grown to early exponential phase in M9-pyr-Cas media. The translational activity of 806 genes was significantly (log_2_>1) affected by *rimBK* deletion (Table S3), with more affected genes showing reduced (564) than increased (242) mRNA abundance in the Δ*rimBK* mutant background. The upregulated *rimBK* regulon was dominated by ribosomal proteins, while the downregulated genes contained a large proportion of ABC amino-acid transporter subunits. We observed a large degree of overlap between this experiment and our published data for the *rimK* proteome (30), with many upregulated genes in the initial (M9-pyr, stationary phase) experiment downregulated in our Ribo-seq dataset and *vice versa* (Table S3).

Our data indicate that RimK produces an indirect, nutrient-dependent effect on the SBW25 translatome by changing the intracellular abundance of Hfq. To test whether RpsF glutamation also exerts a direct effect on SBW25 gene translation, we produced an additional Δ*rimBK* mutant strain in which the *rpsF* gene was extended by 4 glutamate residues. The Δ*rimBK rpsF_4glu_* mutant proved extremely difficult to produce, with the eventual mutant strain displaying a complete loss of UV fluorescent siderophore secretion compared to WT SBW25 (Fig S3). Subsequent whole genome sequencing of this mutant revealed additional SNPs in the *pvdI* and *pvdJ* pyoverdin biosynthetic loci. To investigate the potential significance of these *pvdIJ* mutation we grew SBW25 WT, Δ*rimBK* and Δ*rimBK rpsF_4glu_* strains on KB Congo Red plates containing excess FeCl_2_ to repress pyoverdin production. Under these conditions the Δ*rimBK rpsF_4glu_* mutant displayed enhanced Congo Red binding compared to WT and Δ*rimBK*, suggesting that the *rpsF_4glu_* mutation exerts a *rimBK*/*pvdIJ*-independent effect on SBW25 physiology (Fig S3).

Both Δ*rimBK rpsF_4glu_* and Δ*rimBK rpsF_10glu_* mutants showed a pronounced swarming motility defect compared with WT SBW25 and Δ*rimBK*, while the Δ*rimBK rpsF_4glu_* strain also displayed a significant increase in Congo Red binding. Growth of neither strain was affected in rich (KB) or poor (M9-pyr) media (Fig 10). To investigate the molecular basis of the *rpsF_4glu/10glu_* phenotypes, we conducted Ribo-seq assays for Δ*rimBK rpsF_4glu/10glu_* and compared these data to the translatome of Δ*rimBK*. Translational activity of 159 genes was significantly (>1.0 log_2_) affected by the *rpsF_4glu_* mutation (Fig 11, Table S4), with roughly equal numbers of up- (79) and downregulated (80) genes. Consistent with the observed preference of the RimBK system to produce RpsF_4glu_ ribosomal variants (Fig 4), the *rpsF_10glu_* mutation affected considerably fewer gene targets, with ribosomal activity differing significantly from the Δ*rimBK* background for only 45 loci (Fig S5, Table S4). By comparing the significantly affected genes for Δ*rimBK*/WT against the results for Δ*rimBK rpsF_4glu_*/Δ*rimBK*, we were able to identify three distinct subgroups of loci within the *rpsF_10glu/4glu_* translatomes. The first of these were genes whose translation is altered by *rimBK* deletion, but not by RpsF glutamation (Class 1, Fig 11B). These genes comprised the majority of the *rimBK* regulon, and overlapped substantially with our previously published Hfq translatome dataset (18). Translation of the second set of genes (Class 2, Fig 11C) was shifted by RpsF glutamation and *rimBK* deletion in the same direction (usually down-regulated). This set contained numerous small, uncharacterised proteins alongside several stress response proteins and amino-acid ABC transporters/ metabolic genes (Table S5). We observed a large degree of overlap with the equivalent dataset for RpsfF_10glu_, suggesting that both mutations produce similar effects on this gene class.

**Figure 10:**
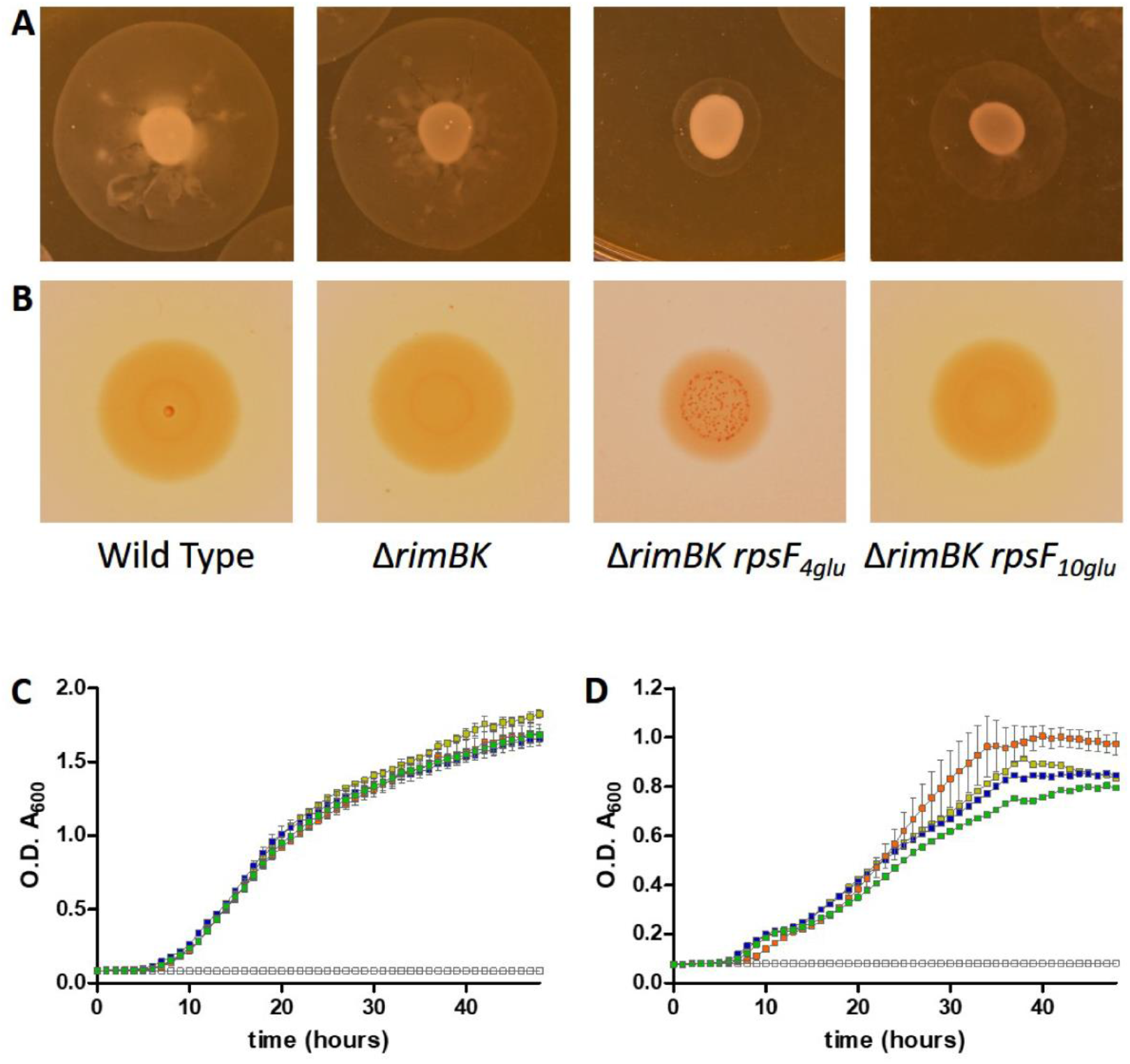
SBW25 *rimBK/rpsF* mutations show distinct phenotypic changes. **(A)** Glutamation of RpsF at the C-terminus influences swarming motility. Overnight growth on swarm diameter plates containing 0.3% w/v agar and 0.05% Congo Red dye. **(B)** Colony morphology resulting from overnight growth of 5 µL spots of the indicated strains, on KB agar plates containing 0.004% Congo Red dye. The presence of four C-terminal glutamates on RpsF results in smaller colonies and increased dye binding. **(C)** Growth curves in KB medium and **(D)** in M9-pyruvate medium. In both charts, WT SBW25 is shown in gold, Δ*rimBK* in blue, Δ*rimBK rpsF_4glu_* in orange and Δ*rimBK rpsF_10glu_* in green. The black line shows the absorbance of the uninoculated media.

**Figure 11:**
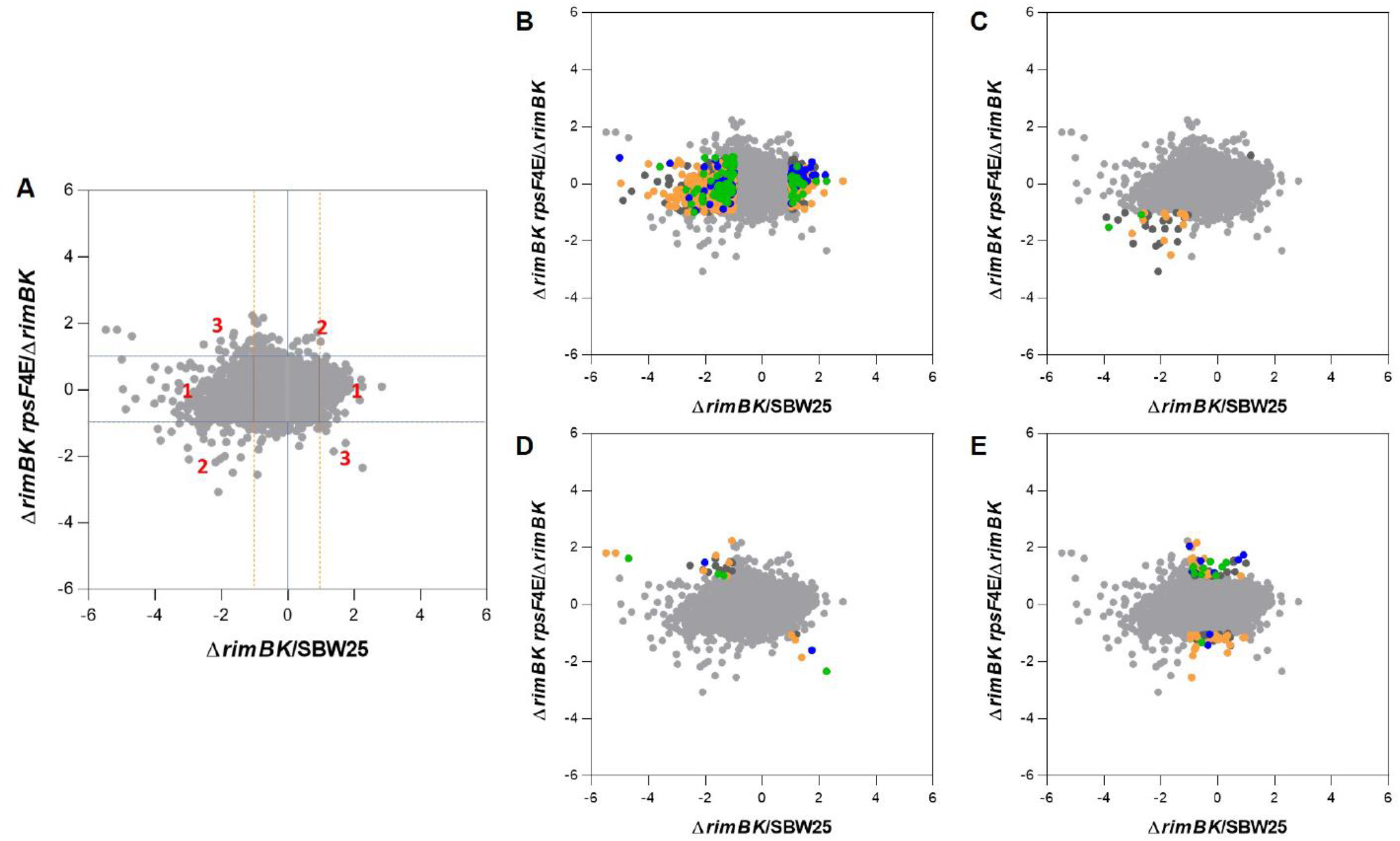
Scatterplots representing the pairwise comparisons of log_2_ ratios between the Δ*rimBK rpsF_4glu_* and Δ*rimBK* translatomes. **(A)** Location of class 1, class 2 and class 3 genes are indicated. Horizontal and orange dashed vertical lines indicate regions of >1.0 log_2_ deviation from zero. **(B-D)** Highlighted regions containing class 1, 2 and 3 genes respectively. **(E)** Genes significantly (>1.0 log_2_) affected by *rpsF* glutamation but not significantly affected by *rimBK* deletion. Highlighted genes are listed in Table S4 and colour-coded according to their COG classifications: yellow = metabolism; green = cellular processes and signalling; blue = information storage and processing; dark grey = poorly characterised.

The third set of genes (Class 3, Fig 11D); those inversely affected by RpsF glutamation and *rimBK* deletion, are candidates for the direct RpsF-glutamation regulon. For RpsF_4glu_, these genes predominantly split into four main classes (Table S5): ABC transporters for fatty/amino-acids, metabolic pathways for metabolism of amino acids, and genes associated with the initial stages of attachment to surfaces (e.g. pili and curli biosynthesis loci) were upregulated by glutamation. As predicted based on the *pvdIJ* mutation, downregulated loci also included the ferripyoverdine receptor *fpvA* (42). RimBK regulation of these key groups (fatty/amino acid transport and metabolism, initial surface attachment and secretion of macromolecules) was also seen when we extended our analysis to include genes that were less than 2-fold different in Δ*rimBK* versus WT, but more than 2-fold inversely affected in *rpsF_4glu_* (Table S5, Figure 11E). This group contained loci for pectate lyase and phytase secretion (both upregulated by glutamation), alongside loci linked to regulation of motility and surface attachment (43). We saw little overlap between the RpsfF_4glu_ and RpsfF_10glu_ regulons for this group of genes, with the smaller, less well-defined 10E regulon containing a few ABC transporter components alongside several hypothetical proteins (Table S5).

## Discussion

Bacterial adaptation to complex natural environments is controlled by an intricate series of connected signalling pathways that function both within individual cells and on the microbial community as a whole (22, 25, 44). In addition to extensive transcriptional (17, 22) and protein functional regulation (23), control of mRNA translation is critical for effective colonisation of plant rhizospheres (18). In this study we characterise the RimABK pathway in *Pseudomonas fluorescens*; a novel translational regulatory system that controls bacterial adaptation to the rhizosphere environment through specific, controlled modification of a ribosomal protein.

The activity of the RimK glutamate ligase is controlled by several distinct environmental inputs and the intracellular levels of key molecules. At the transcriptional level, expression of the polycistronic *rimABK* mRNA is controlled by temperature and nutrient availability, with cold, nutrient-starved conditions leading to increased *rim* transcript abundance. At this stage the transcriptional regulators that control *rim* expression are unknown. The environmental cues that activate *rim* transcription would make the operon a plausible target for σ^S^ regulation, although this was not supported by a recent study of sigma-factor regulation in *P. aeruginosa* (45). A further, substantial fraction of RimK regulation occurs at the protein activity level, where RimA, RimB and intracellular levels of glutamate and cdG combine to translate a complex set of environmental variables into a single output: the proportion of all ribosomally-associated RpsF proteins that have C-terminal poly-glutamate tails.

Our results suggest that in the presence of RpsF, RimK adds glutamate units and RimB removes them. The length of poly-glutamate tails is restricted, tending towards a minimal number of 4, as a consequence of the specific protease activity of RimB. With low levels of cdG in the system, a balance is established where RimK activity predominates and a range of chain lengths (longer than four) arises. This may also reduce the fraction of RpsF proteins that can be glutamated. In the presence of RimB and with increasing cdG levels, RimA activity switches from its default role of stimulating RimK to degrading cdG, leading to a reduction in RimK activity and an increased ratio of RimB to RimK activity (Fig S6). RimA thus functions as a cdG-trigger enzyme (46): a signalling protein whose enzymatic activity changes its interaction with a regulatory partner, in this case RimK. The data showing that both RimA and cdG increase RimK ATPase and glutamate ligase activity when no RimB is in the system, suggests that this trigger behaviour of RimA requires of presence of RimB. The kinetic details of this mechanism remain unclear.

The cdG-induced switch of RimA activity results in shortened RpsF chain lengths and higher overall coverage. RimA phosphodiesterase activity may also serve to dampen the effect of transient fluctuations in cdG level on RimK activity. Short impulses of cdG would be rapidly degraded by RimA and RimK activity would quickly return to the default position of making longer chains. Only if cdG is produced and present in sufficient quantities for a defined time would RimK glutamation activity drop, thus enabling RimB to limit RpsF chain lengths and move towards global RpsF coverage. The trade-off between long chains and high RpsF coverage is therefore determined by cdG availability. To enable RpsF modification to function as an effective regulator, a mechanism should also exist to return modified RpsF to a non-glutamated ‘ground state’ (Fig 7). Our conceptual model for Rim function is summarised in Fig 12.

**Figure 12:**
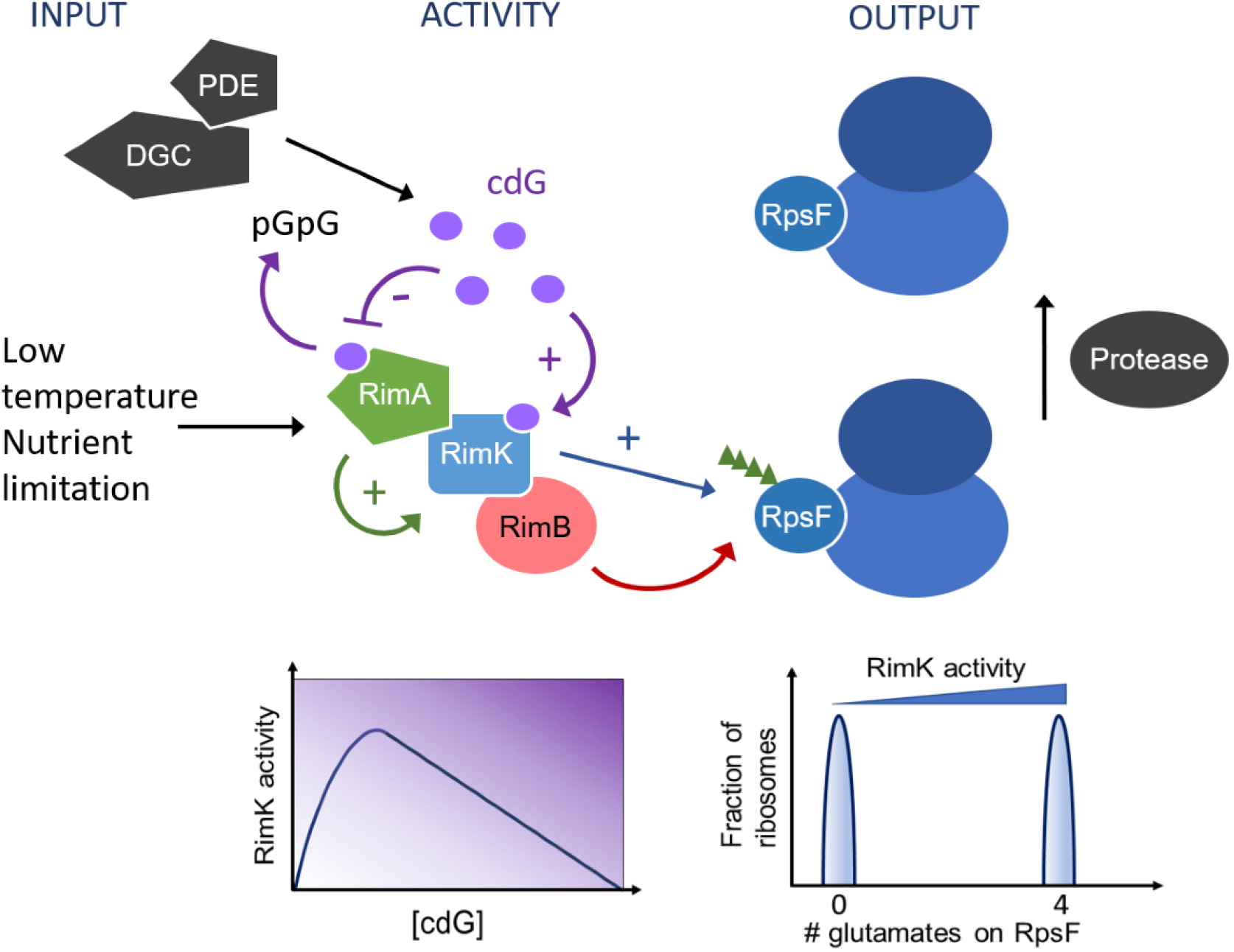
A model for RimABK/RpsF regulation in *P. fluorescens*. Activity of RimK (blue rectangle) is stimulated by direct interaction with RimA (green pentangle) and cdG (purple circles). CdG also interacts with RimA, leading to its hydrolysis to the linear dinucleotide pGpG and an accompanying loss of RimA stimulation. The combined activity of the glutamate ligase RimK and the poly-glutamate protease RimB (red oval) adds chains of ∼4 glutamate residues (green triangles) to the C-terminus of ribosomal protein RpsF (blue oval). Known inputs to the system include changes in *rimABK* expression, stimulated by low temperature and nutrient limitation, glutamate concentration, and cdG concentration. CdG levels are controlled in turn by as-yet-unidentified diguanylate cyclases/phosphodiesterases (grey pentangles). These inputs combine to control the proportion of glutamated ribosomes in the cell. Glutamated ribosomes are returned to a non-glutamated state by the action of an as-yet-unidentified protease (grey oval). Insets below the main diagram illustrate the impact of increasing cdG concentration on RimK activity (left, Fig 3, Fig S5) and the predicted relationship between RimK activity and ribosomal glutamation (right, Fig 6).

The proportion of modified versus unmodified RpsF proteins leads to altered proteome composition and translational activity, which can be effectively measured using Ribo-seq. This sequencing-led approach measures global ribosomal activity towards different mRNAs. We have previously shown that Ribo-seq provides a more accurate assessment of translational regulation than either RNA-seq, which does not record translation level control, or quantitative proteomic analyses that are subject to extensive, compensatory post-translational effects (18). The implications of Rim activity are substantial, with the translation of several hundred genes significantly affected in a Δ*rimK* mutant strain.

Much of the altered translational activity in Δ*rimK* can be explained by changes in Hfq abundance. Consistent with this, the mobile tails of RpsF and the neighbouring ribosomal protein S18 have been shown to stochastically interact with protein S1 in the *E. coli* ribosome (47). Protein S1 is required for translation initiation of mRNAs with structured 5’ ends or with weak or no Shine-Dalgarno sequences (48). Furthermore, in *E. coli* S1 weakly and reversibly associates with the ribosome and interacts with Hfq when binding RNA polymerase to affect transcription (49). In *P. aeruginosa*, Hfq interaction with nascent transcripts has been proposed as a mechanism for controlling translation in cases where transcription and translation are coupled (50). Additionally, Hfq has been shown to associate with *E. coli* ribosomes, and both Hfq and S1 are components of the bacteriophage Qβ replication complex (51–53). It is possible therefore that modification of RpsF by C-terminal glutamation may disturb the binding equilibrium between protein S1 and the ribosome, influencing the cellular localisation of S1 and Hfq. A redistribution of S1 and Hfq within the cell would be expected to exert a significant effect on protein abundance and may explain a substantial fraction of the observed RimK regulatory activity.

While RimK disruption or RpsF modification both affect the *P. fluorescens* proteome, the addition of four glutamate residues to RpsF has additional biological significance. Strains with the *rpsF_4glu_* allele display a shift in translational activity for a set of genes associated with early-stage surface attachment, amino-acid utilisation and transport and the secretion of molecules (e.g. phytase and pectate lyase) associated with rhizosphere colonisation. RpsF modification specifically affects translation and proteome composition within minutes of detecting an environmental shift. This strongly suggests that glutamation by RimK is not a passive process determining long-term population level adaptation, but rather is an active mechanism used by individual cells to rapidly adapt to changes in their environments by modifying mRNA translation activity.

We are now able to propose a general mechanism for RimK translational control (Fig 12), Nonetheless, several key questions remain to be answered. Firstly, the exact mechanism of translational discrimination by ribosomes containing modified and unmodified RpsF is still unclear. We were unable to identify any clear distinguishing features in the primary or predicted secondary structures of the Rim-mRNAs, nor could we assign them definitively to a single transcriptional or translational regulon. Given the highly pleiotropic effects of *rim* gene disruption, the mechanism of RpsF-discrimination likely awaits a comparative structural analysis of modified and unmodified *Pseudomonas* ribosomes. More generally, the identity of several important pathway components; specifically, the ‘reset’ protease that removes 4E tails from RpsF, and the DGCs/PDEs that control the cdG input to RimA/RimK remain to be determined. These proteins present additional opportunities for regulatory input to the Rim pathway, increasing the complexity of the system and further integrating it into the signalling network that controls *Pseudomonas* behaviour.

Finally, a comparative examination of the Rim proteins from different bacterial species may provide interesting new insights into the function and biological significance of the RimABK pathway and its individual component proteins. RimK homologs have been linked to stress-tolerance and pathogenicity in several human and plant pathogens (30, 54), although the phenotypic consequences of *rimK* deletion vary widely between different organisms (30). Furthermore, variations on the classical three-gene Rim pathway are widespread in nature, with RimBK, RimAK and RimK-only variants identified in diverse bacterial families. Homologs of *rimK* have also been identified in several eukaryotic genomes. Overall, our study suggests that dynamic, post-translational ribosomal modification is a common feature of prokaryotic, and potentially archaeal and eukaryotic environmental adaptation.

## Materials and Methods

### Strains and growth conditions

Plasmids and strains are listed in Table S6. Primers are listed in Table S7. Unless otherwise stated all *P. fluorescens* strains were grown at 28°C and *E. coli* at 37°C in lysogenic broth (LB) (55) solidified with 1.3% agar where appropriate. Gentamycin was used at 25 μg/ml, carbenicillin at 100 μg/ml, and tetracycline (Tet) at 12.5 μg/ml. For inducible plasmids, IPTG was added to a final concentration 0.2 mM (SBW25) or 1 mM (*E. coli*), unless otherwise stated.

### Molecular microbiology procedures

Cloning was carried out in accordance with standard molecular microbiology techniques. The pETM11-*rpsF_10glu_*, pETM11-*rimA_E47A_* and *E. coli rpsF* purification vectors were produced by ligating PCR fragments (amplified with primers 17/18, 11/12/15/16 and 13/14 respectively) between the *Nde*I and *Xho*I sites of plasmid pET*Nde*M-11 (56) as appropriate. Purification vectors for *rimA, rimB* and *rimK* were as described in (30) using primers 7-12. *P. fluorescens* SBW25 site-specific mutants were constructed using a modification of the protocol described elsewhere (57). Up- and downstream flanking regions to the target modification site (*rimBK*, *hfq* or *rpsF*) were amplified using primers 19-25, 26-31 and 32-37 respectively. PCR products in each case were ligated into pTS1 between *Kpn*I/*EcoR*I, *Nde*I/*BamH*I and *Xho*I/*Kpn*I respectively. The resulting vectors were transformed into the target strain, and single crossovers were selected on Tet and re-streaked for single colonies. Cultures from single crossovers were grown overnight in LB medium, then a dilution series plated onto LB plates containing 5% w/v sucrose to enable *sacB* counterselection. Individual sucrose-resistant colonies were patched onto LB plates ± Tet, and Tet-sensitive colonies tested for successful gene modification by colony PCR followed by Sanger sequencing. Reverse transcriptase PCR (RT-PCR) and 5’ RACE were conducted using primers (1/2 and 3/4 respectively). qRT-PCR was performed as described in (30) using primers 5/6. qRT-PCR experiments were repeated three times independently and data are presented +/− standard error.

### Purification of Rim and RpsF proteins

Protein purification was conducted after (30). Briefly, overnight cultures of *E. coli* BL21-(DE3) pLysS were used to inoculate 1.0 litre overexpression cultures. These were then grown at 30 °C to an OD_600_ of 0.6, before protein expression was induced for 2 hours with 1mM IPTG. Cells were lysed using a cell disruptor (Avestin) and His_6_-tagged proteins purified by NTA-Ni chromatography. 1 ml HiTrap chelating HP columns (Amersham) were equilibrated with 25 mM KH_2_PO_4_, 200 mM NaCl, pH 8.0 (SBW25 RimA/B/K), 50 mM Tris-Cl, 2.5% glycerol, pH 8.0 (SBW25 RpsF) or 50 mM Tris, 300 mM NaCl, 10 mM imidazole, pH 9.0 (*E. coli* RpsF), and loaded with cell lysate. Following protein immobilization, proteins were eluted with a linear gradient to 500 mM imidazole over a 15 ml elution volume.

### Linked Pyruvate Kinase / Lactate Dehydrogenase (PK/LDH) ATPase activity assays

ATPase activity was measured indirectly by monitoring NADH oxidation in a microplate spectrophotometer (BioTek Instruments) at 25 °C. The reaction buffer consisted of 100 mM Tris-Cl (pH 9.0), 20 mM MgSO_4_. Each 100 μL reaction contained 0.4 mM NADH, 0.8 mM phosphoenolpyruvic acid, 0.7 μL PK/LDH (Sigma) and was initiated by the addition of 10 μL ATP. Protein concentrations were as shown in the figure legends. Enzyme kinetics were determined by measuring A_340_ at 1-minute intervals. Kinetic parameters were calculated by plotting the specific activity of the enzyme (nmol ATP hydrolyzed/min/mg protein) versus ATP concentration and by fitting the non-linear enzyme kinetics model (Michaelis-Menten) in GraphPad Prism. 25 μM cyclic dinucleotides were included where appropriate, as noted in the text.

### RpsF glutamation and protease assays

The glutamation assay was adapted from (40). Briefly, purified RpsF and Rim proteins at the concentrations indicated in the figure legend were incubated at room temperature for the indicated times in reaction buffer comprising 100 mM Tris-HCl pH 9.0, 20 mM L-glutamate (unless otherwise indicated), 20 mM ATP and 20 mM MgSO_4_·7H_2_O. Reactions were supplemented with cdG (Sigma) as indicated in the text. Protease assays were performed in 100 mM TRIS-HCl pH 9.0, 20 mM L-glutamate, 20 mM ATP, 20 mM MgSO_4_·7H_2_O with RimB concentrations as stated in the figure legend. For both experiments, samples were boiled in SDS loading buffer and analyzed by SDS-PAGE following completion of the reaction.

### Growth Assays

Bacterial growth was measured in a microplate spectrometer (BMG Labtech FLUOstar Omega) using 2 biological duplicates each providing 2 experimental replicates and are presented as mean +/− standard error. 150 µL of the indicated growth medium in each case was added to the internal 60 wells of clear-bottomed, black-walled 96-well microplates (Sigma). Carbon sources were added to a concentration of 0.4% w/v in each case. Growth was initiated by the addition of 5 µL of overnight cell culture (LB media, 28 °C, shaking) normalized to an OD_600_ of 0.01. Plates were incubated statically at 28 °C and agitated immediately prior to each data acquisition step. Optical density was measured at 600nm and fluorescence was monitored at 460nm with an excitation wavelength of 355nm. Experiments were conducted at least twice independently.

### Phenotypic assays

To assess the colony morphology of the SBW25 *rim/rpsF* mutants, Kings B (KB) plates (58) were prepared containing 0.8% w/v agar, 1% w/v NaCl, 0.004% Congo Red dye, +/− 100 µM FeCl_2_. Plates were allowed to dry for 45 min in a sterile flow chamber. Meanwhile SBW25 strains were grown over-day at 28°C in LB media with shaking. Cell densities were adjusted to give an optical density (λ 600nm) of 0.2 then 1.5 µL of each culture was spotted onto the media surface. Six replicates were prepared from two biological duplicates (three plates prepared from each). Plates were then incubated at 28 °C for the time indicated in the figure legends before photographing, with a representative image shown in each case.

To quantify swarming diameter, Kings B (KB) media plates containing 0.3% agar w/v, 1% w/v NaCl, 0.004% Congo Red dye were produced and allowed to set and dry for 45 min in a sterile flow chamber. SBW25 strains were grown overnight at 28°C in LB media with shaking. The following morning, cell densities were adjusted to an optical density (λ 600nm) of 0.2 then 5 µL of each culture was spotted onto the media surface. Six replicates were prepared from two biological duplicates (three plates prepared from each). Plates were then incubated over-day at 28 °C then overnight at 4 °C before photographing, with a representative image shown in each case. All phenotypic experiments were conducted at least twice independently.

### Determining the transcription start site of the *rim* operon

5’ RACE (Life Technologies) was performed on mRNA prepared from SBW25 cells grown overnight in M9 media with pyruvate (0.4% w/v) as sole carbon source. RNA was purified by column capture (Qiagen RNeasy Mini Kit). Purified RNA was subjected to additional DNase treatment (Turbo™ DNase, Ambion). The final PCR product from the 5’ RACE procedure was excised from a 1% agarose gel and purified by column capture (Macherey-Nagel Nucleospin mini kit). The purified product was then Sanger sequenced (Eurofins Scientific) to reveal the position of the transcriptional start site.

### RT-PCR

cDNA from SBW25 cells was prepared as an independent step prior to PCR. Total RNA was extracted from cells grown in M9 media containing pyruvate (0.4% w/v) as sole carbon source by column capture (Qiagen RNeasy Mini Kit). Purified RNA was subjected to additional DNase treatment (Turbo™ DNase, Ambion). The resulting RNA was then subjected to amplification (Merck WTA2 Amplification Kit) to produce cDNA that was subsequently purified by column capture (Qiagen QiaQuick PCR purification kit) and eluted in RNase-free water. PCR was performed on the cDNA using high-fidelity DNA polymerase (New England Biolabs Phusion polymerase) using a forward primer targeting the start of *rimA* (*PFLU0263*) and a reverse primer targeting the end of *rimK* (*PFLU0261*). The resulting PCR product was purified by column capture (Macherey-Nagel Nucleospin mini kit) and sequenced (Eurofins Scientific) to confirm the presence of the three *rim* genes.

### Automated Immunodetection

Biological duplicate SBW25 cultures were grown overnight in LB media at 28 °C. These cultures were used to inoculate 50ml volumes of M9-pyruvate minimal media, with and without Cas-aminoacids. Pyruvate and Cas-aminoacids were used at a concentration of 0.4% w/v. When cultures reached an optical density of 0.3 (measured at 600 nm), 1 ml of culture was removed, centrifuged and the resulting pellet resuspended into 100 µL supernatant + 100 µL x2 SDS loading buffer. The remaining cultures were allowed to grow overnight to reach stationary phase. 150 µL of a WT culture grown in M9-Pyr-CAS was taken and a proportionate volume of all other cultures (according to differences in optical density measured at 600nm) was taken. Pellets were resuspended as above. Prior to immunodetection, samples were briefly sonicated to reduce viscosity. 0.5 µL of each sample was run on WES™ ProteinSimple automated immunodetection system (Bio-Techne) according to the manufacturer’s instructions.

### Western Blotting

Following growth in M9-Pyr-CAS, the concentration of the soluble protein fraction from SBW25 cells was normalized. 140µg total protein from each sample was loaded in a 25µL volume onto one of two SDS-PAGE gels. Gels were blotted onto polyvinylidene difluoride (PVDF) membranes (Millipore). Membranes were incubated overnight in blocking solution (1x PBS pH 7.4, 0.01% Tween 20, 5% milk powder). Protein was subsequently detected with either 1/2000 anti-polyglutamate chain (AdipoGen) or 1/200 anti-RpsF (Dundee Cell Products) antibodies and 1/6000 anti-rabbit secondary antibody (Sigma). Bound antibody was visualized using ECL chemiluminescent detection reagent (GE Healthcare).

### Protein Extraction and Mass Spectrometry

SDS gel slices containing protein samples of interest were washed, treated with DTT and iodoacetamide, and digested with trypsin according to standard procedures. Peptides were extracted from the gels and analysed by LC-MS/MS on an Orbitrap-Fusion™ mass spectrometer (Thermo Fisher, Hemel Hempstead, UK) equipped with an UltiMate™ 3000 RSLCnano System using an Acclaim PepMap C18 column (2 µm, 75 µm x 500mm, Thermo). Aliquots of the tryptic digests were loaded and trapped using a pre-column which was then switched in-line to the analytical column for separation. Peptides were separated using a gradient of acetonitrile at a flow rate of 0.25 µl min-1 with the following steps of solvents A (water, 0.1% formic acid) and B (80% acetonitrile, 0.15 formic acid): 0-3 min 3% B (trap only); 3-4 min increase B to 7%; 5-40 min increase B to 50%; 40-45 min increase B to 65%; followed by a ramp to 99% B and re-equilibration to 3% B.

Data dependent analysis was performed using HCD fragmentation with the following parameters: positive ion mode, orbitrap MS resolution = 120k, mass range (quadrupole) = 300-1800 m/z, MS2 top20 in ion trap, threshold 1.5e4, isolation window 1.6 Da, charge states 2-7, AGC target 2e4, max inject time 35 ms, dynamic exclusion 1 count, 15 s exclusion, exclusion mass window ±5 ppm. MS scans were saved in profile mode and MS2 scans were saved in centroid mode.

To generate recalibrated peaklists, MaxQuant 1.6.0.16 was used, and the database search was performed with the generated HCD peak lists using Mascot 2.4.1 (Matrixscience, UK). The search was performed on a *Pseudomonas fluorescens* SBW25 protein database (Uniprot, 20140727, 21,935 sequences) to which copies of the RpsF sequence with 1-10 additional glutamate residues (E) at the C-terminus was added. For the search a precursor tolerance of 6 ppm and a fragment tolerance of 0.6 Da was used. The enzyme was set to trypsin/P with a maximum of 2 allowed missed cleavages. Oxidation (M), deamidation (N, Q), and acetylation (N-terminus) were set as variable modifications, carbamido-methylation (CAM) of cysteine as fixed modification. The Mascot search results were imported into Scaffold 4.4.1.1 (www.proteomsoftware.com) using identification probabilities of 99% for proteins and 95% or 0% for peptides, as discussed in the results.

### Quantitative Mass Spectrometric Analysis Using Isobaric Labelling (iTRAQ)

50 ml overnight cultures of SBW25 WT and Δ*rimK* were grown at 28 °C in LB medium with shaking. Cultures were then pelleted by centrifugation and resuspended in equal volumes of A) carbon-free rooting solution (30), pre-cooled to 8 °C, and B) LB medium, pre-heated to 28 °C. Cultures were incubated for 45 minutes with shaking at 8 or 28 °C as appropriate, then cellular activity was halted by addition of 30 ml of RNAlater [saturated (NH_4_)_2_SO_4_, 16.7 mM Na-Citrate, 13.3 mM EDTA, pH 5.2] containing protease inhibitors. Cell samples were then pelleted, washed three times with 10 mM HEPES pH 8.0 + protease inhibitors, before re-suspension to a final volume of 200 μL. 700 μL pre-cooled RLT + β-mercaptoethanol buffer (RNeasy Mini Kit, QIAGEN) was added and samples lysed with two 30 s Ribolyser pulses at speed setting 6.5. Supernatant was removed, and the soluble fractions separated by ultracentrifugation (279,000 *g*, 30 min, 4 °C).

After determination of protein concentration, the soluble proteins were precipitated with chloroform-methanol and subjected to iTRAQ 4-plex quantification. The experiment was performed with 2 biological replicates. In each replicate the samples were labelled with an iTRAQ 4 plex kit (Sciex) as follows: 114=WT LB28, 115= Δ*rimK* LB28, 116=WT RS8, 117= Δ*rimK* RS8. The labelled samples were combined and fractionated using the Pierce™ High pH Reversed-Phase Peptide Fractionation Kit producing 7-8 fractions. After analysis on an Orbitrap Fusion (Thermo) using MS3 synchronous precursor selection based on methods described previously (18), the raw data from both replicates were combined and processed in Proteome Discoverer 2.4 (Thermo) with the following main parameters: protein sequence database: *P. fluorescens* SBW25 (Uniprot, Feb/2016, 6388 entries); variable modifications: oxidation (M), deamidation (N,Q); Percolator strict FDR target: 0.01; reporter ion quantifier: most confident centroid, 20 ppm, HCD, MS3; consensus workflow for statistical analysis: replicates with nested design; use unique peptides for protein groups only, co-isolation threshold 50%, imputation: low abundance resampling, ratio based on pairwise ratios, hypothesis test: t-test (background based) generating adjusted p-values according to Benjamini-Hochberg. The protein results table was exported from Proteome Discoverer and used to generate the final protein expression tables and plots (Fig 8, Fig S4, Table S2) in The R Project for Statistical Computing.

### Ribosomal profiling (Ribo-Seq) analysis

Ribosomal profiling was conducted as described in (18). SBW25 cultures were grown at 28°C in M9-pyr-CAS medium to the late exponential phase, then cells were harvested by rapid filtration as described in (59). Collected cells were flash frozen in liquid nitrogen and cryogenically pulverized by mixer milling (Retsch), then thawed and clarified by centrifugation. Resulting lysates were digested with MNase, quenched with EGTA and resolved by sucrose density gradient ultracentrifugation. Ribosome-protected mRNA footprints were processed as previously described (18, 59) and sequenced by Illumina HiSeq2000. Sequencing reads in fastq files were adaptor trimmed using a Perl script that implemented the procedure described in (60). Next, ribosomal RNA sequences were filtered out by aligning them against a Bowtie2 index of the SBW25 rRNAs. The remaining reads were then aligned to the SBW25 genome to produce SAM files, which were used to calculate the centre-weighted coverage at each nucleotide position in the genome. For this, we used a Perl script to select alignments of between 23 and 41 nucleotides in length and counted for nucleotide positions after trimming 11 nucleotide positions from either end of the alignment. This was done separately for reads aligning to the forward and reverse genome strands and the centre-weighted coverage was stored in separate files for each strand. A separate Perl script was used to calculate RPKM values for each gene based on strand specific centre-weighted coverages along the genome. The limma function plotMDS was then used to make PCA plots. The ribosome profiling data has been submitted to ArrayExpress, with accession number E-MTAB-5408.

### Kinetic modelling

For each model, the chemical equations were translated into ordinary differential equations using mass action kinetics (Table S1). Thermodynamic constraints were considered to ensure consistency between reaction rates via different routes, thereby reducing the number of free kinetic parameters. Kinetic forward rate constants were set to 10^9^ M^−1^s^−1^ (diffusion limited) and equilibrium dissociation constants, K_d_, were – based on previous estimates – set initially to 1 µM and then optimized to fit with experimental data (see below). We used the ODE solver Vern7 from the Julia Differential Equations library to solve the chemical kinetics equations, which is suitable for high accuracy, non-stiff systems. Initial concentrations for cdG, RimK, RimA, and RimB were set to 1 µM unless specified otherwise, for instance when mimicking the conditions of a specific experiment as in Fig S2 where [cdG] was 25 µM. Solution of the ODE allowed us to determine steady state values for all RimK species for each model. These steady state concentrations were used to calculate the enzymatic modification of RpsF. K_m_ and k_cat_ values for each species of RimK and K_d_ values between Rim proteins were optimized to fit the model simulations to available experimental data, Figure S2, using the Julia BlackBoxOptim library. Contributions of each RimK species to the kinetics with multiple species present were determined via optimization using all available data set simultaneously using Michaelis-Menton equations for the enzymatic reactions. Glutamation of the population was computed using a stochastic simulation that consisted of the addition of glutamate unit, the removal of a unit, or doing nothing. These probabilities are dependent on the concentration of the various RimK species and RimB. Average behaviour was computed from 1,000,000 independent stochastic simulations. All code was written in the free and open source scientific programming language Julia.

## Acknowledgements

The authors would like to thank Carlo de Oliveira Martins for help and advice with the proteomic analysis. This work was funded by BBSRC Institute Strategic Program Grants BB/J004553/1 (*Biotic Interactions*) and BBS/E/J/000PR9797 (*Plant Health*) to the John Innes Centre, and BBSRC Responsive Mode Grant BB/M002586/1 to JGM.

---

**Figure S1:**
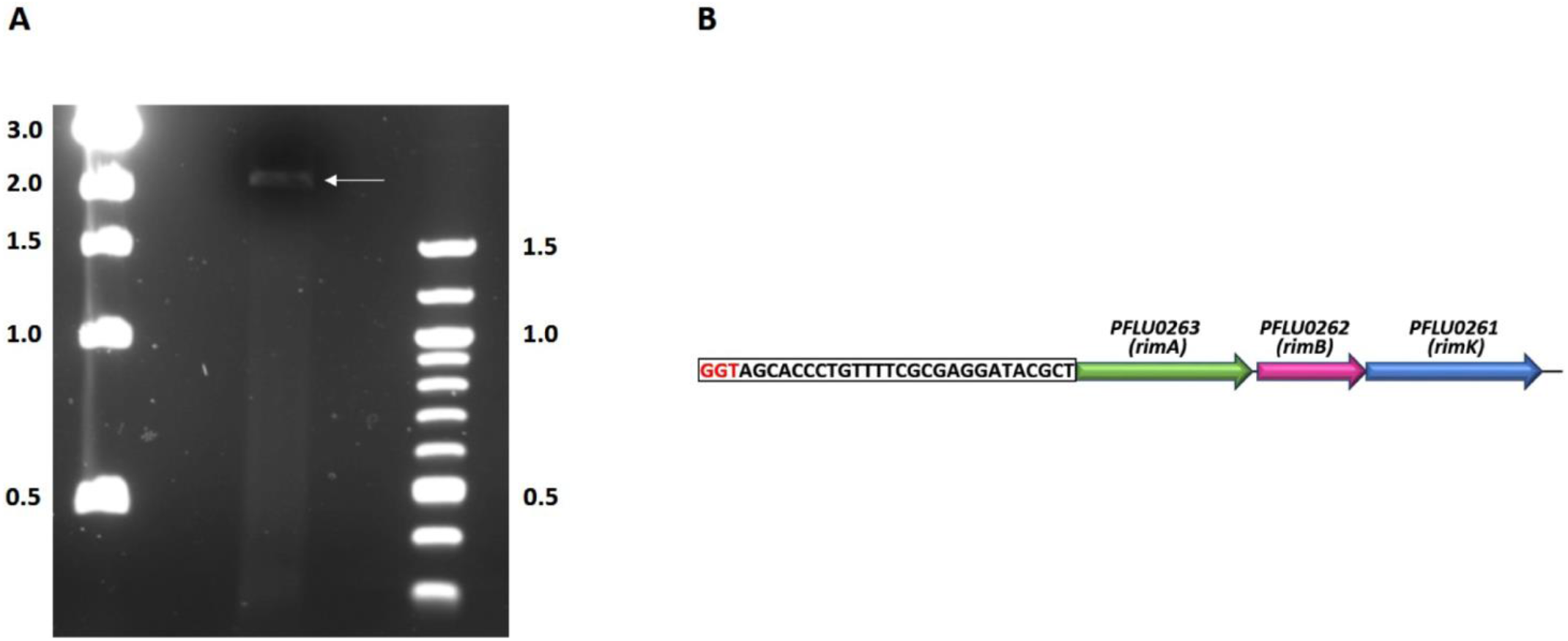
(A) PCR of the *rim* operon from cDNA. The indicated band shows the position of the 2.2Kbp PCR product resulting from the amplification of the *rimABK* operon. **(B) Determining the transcription start site of the *rimABK* operon.** A cartoon representation of the *rimABK* operon and upstream region. Transcription initiation begins at one of the three nucleotides highlighted in red. The 5’ RACE methodology does not allow discrimination between these nucleotides.

**Figure S2:**
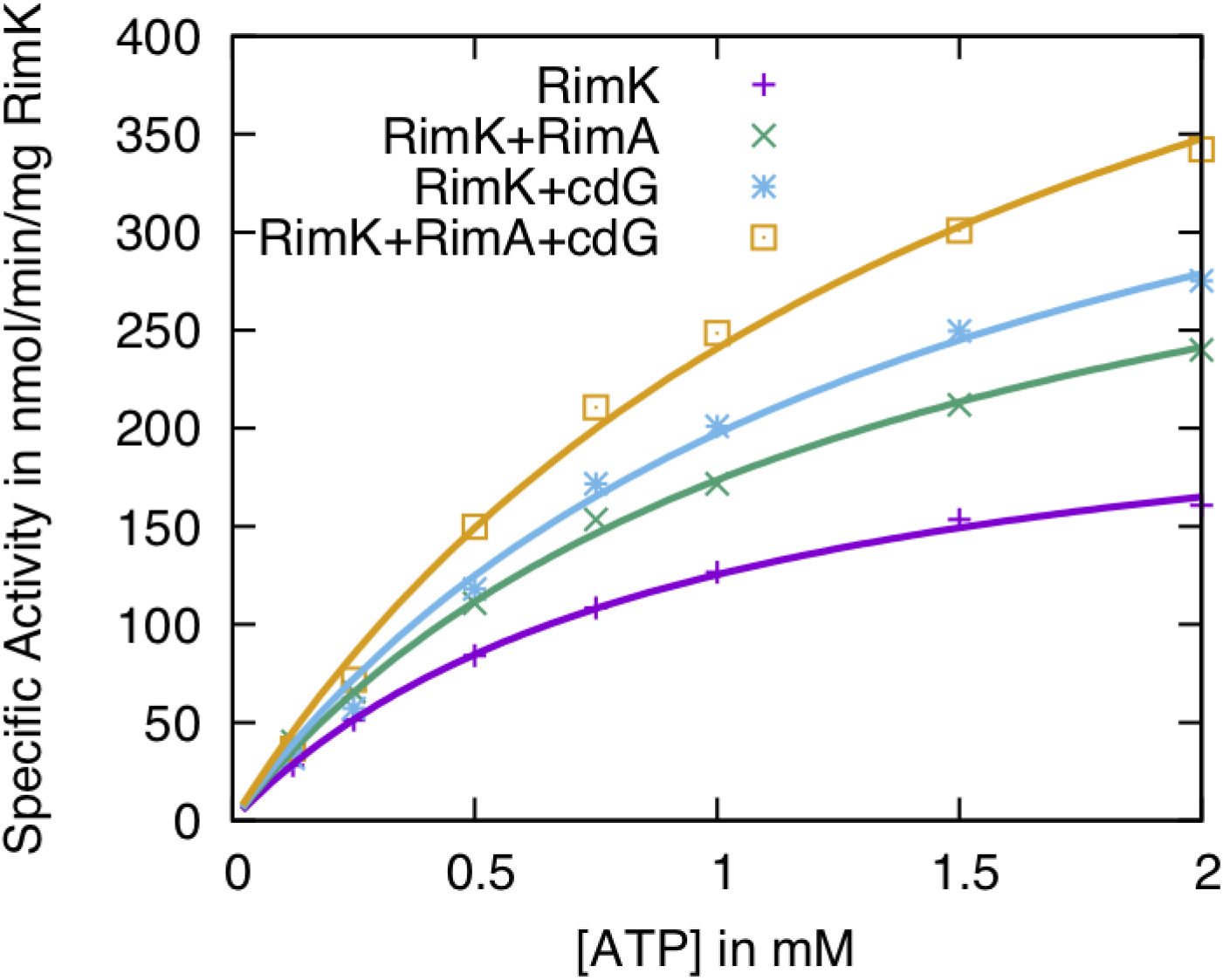
ATPase assays suggest different activation states for RimK. This plot shows the experimental data from Fig 2E as points and the curves are computed from kinetic models based on RimA and cdG binding to RimK (Table S1). Under the assumption that RimK can exist in two states, RimK and RimK*, with different activities and that RimK* achieved either by binding cdG or RimA, we simultaneously optimised all the parameters in the system (percentage of RimK*, k_cat_* and K_m_*). This plot shows that a simple two state model is consistent with the data. However, under this hypothesis and with [RimK]=1 µM, [RimA]= 1 µM, [cdG] = 25 µM, the best fit to the ATPase data is achieved for an equilibrium dissociation constant (K_d_) of 83.2 nM between RimK and RimA and 15.1 µM between RimK and cdG. This it at odds with independent measurements that estimate K_d_ between RimK and cdG to be about 1 µM (30). This suggests that the simple two state system is unlikely and that RimK can exist in at least four different activation states (RimK, RimK.RimA, RimK.cdG and RimK.cdG.RimA, Table S1). This model of RimK ATPase activity results in an insignificantly small better fit to the ATPase activity data but with a K_d_ between RimK and cdG of 1 µM and a K_d_ between RimK and RimA of 0.2 µM.

**Figure S3:**
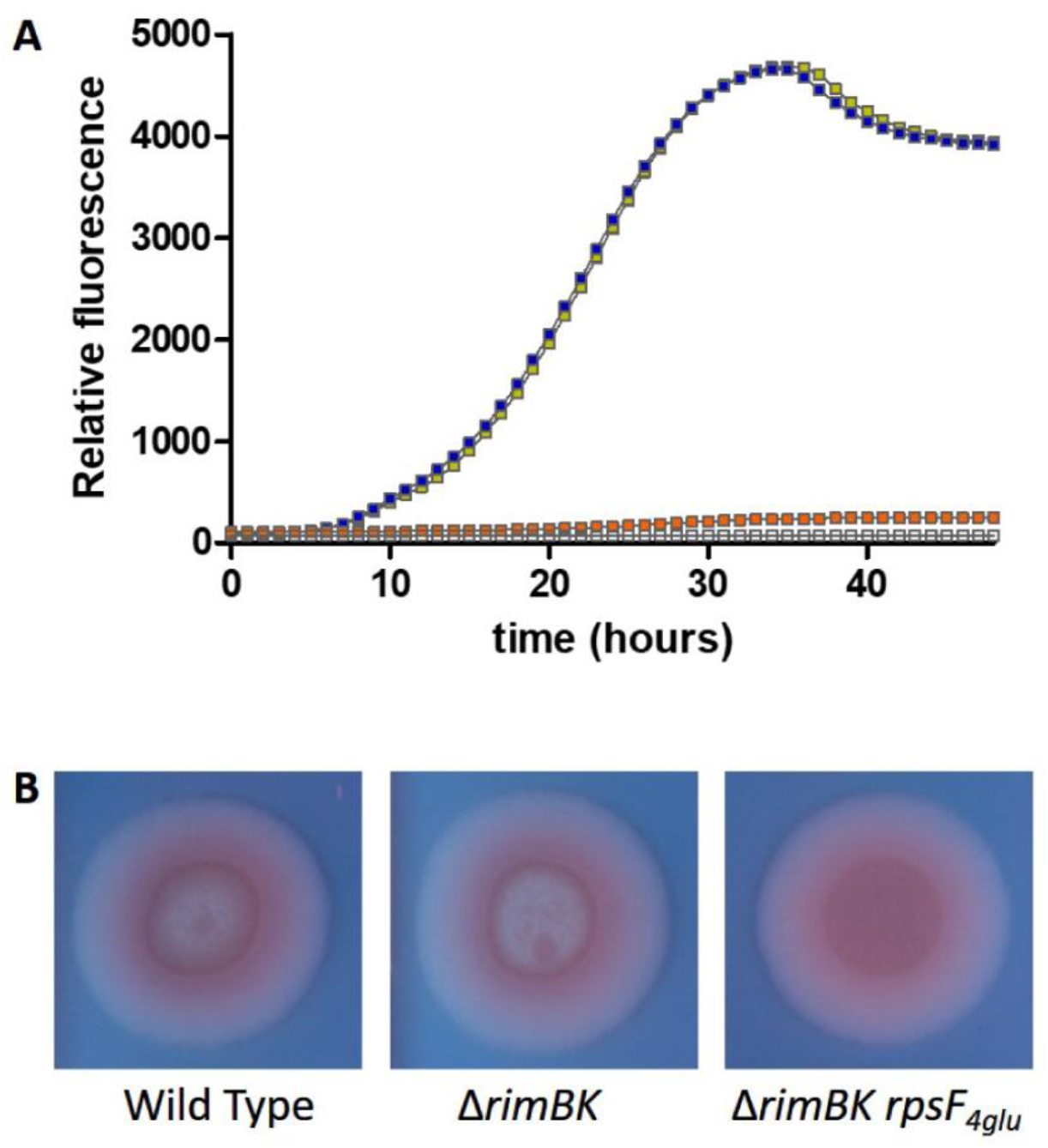
The Δ*rimBK rpsF_4glu_* mutant shows a loss of UV fluorescence and increased Congo Red binding on iron replete media. **(A)** Relative fluorescence (arbitrary units) at A_460_ of SBW25 mutant strains. WT SBW25 is shown in gold, Δ*rimBK* in blue and Δ*rimBK rpsF_4glu_* in orange. **(B)** Colony morphology resulting from 72-hour growth of 5 µL spots of the indicated strains, on M9-pyruvate plates containing 0.004% Congo Red dye and 100 µM FeCl_2_.

**Figure S4:**
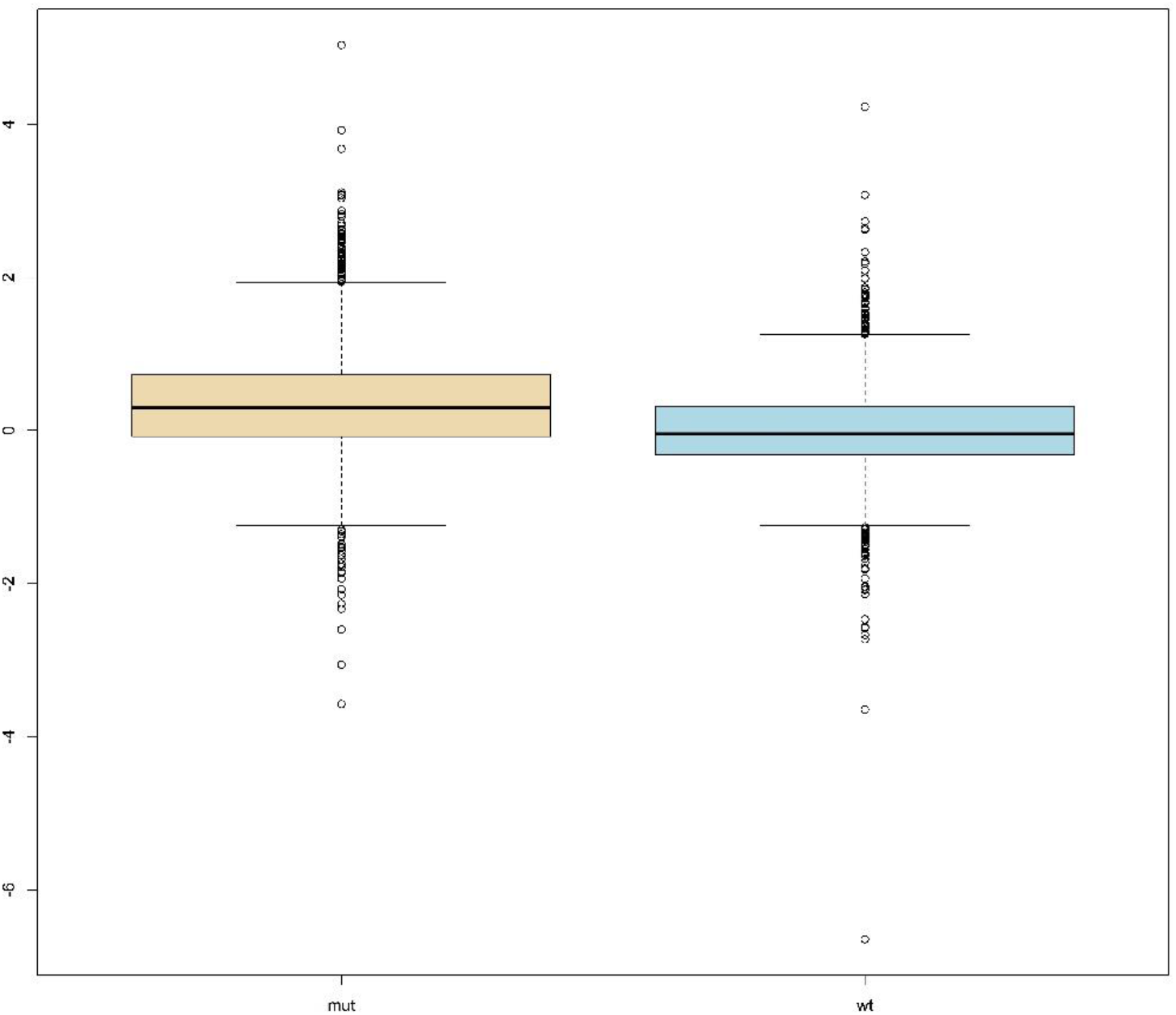
Distribution of all proteins quantified in Proteome Discoverer software (see iTRAQ method section) upon transition to cold, nutrient limiting conditions. Boxplots showing log_2_-fold change ratios for protein abundance after: [45 minutes incubation in carbon-free rooting solution at 8 °C]/[45 minutes incubation in LB medium at 28 °C], for Δ*rimK* (mut) and WT SBW25 (wt).

**Figure S5:**
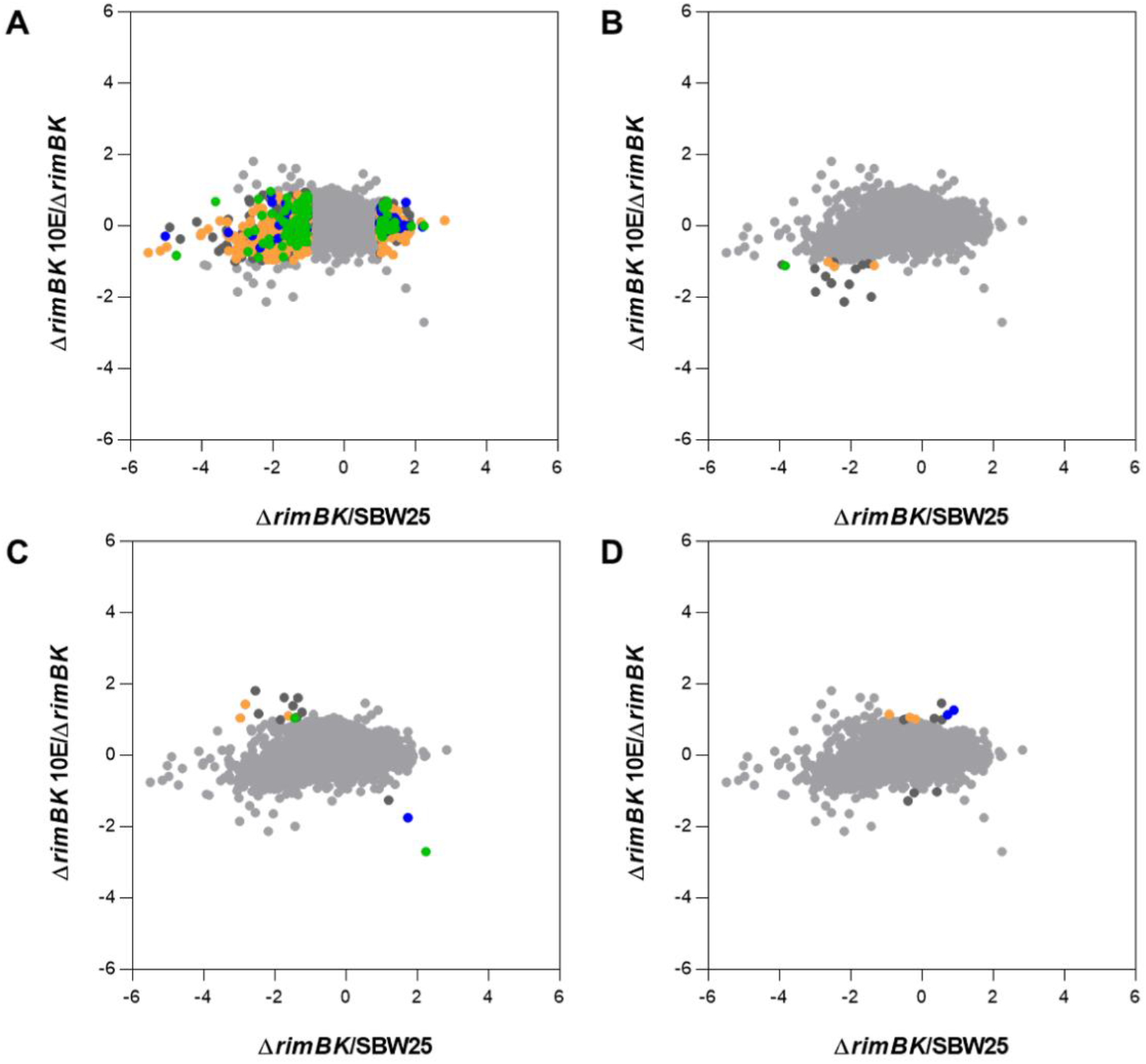
Scatterplots representing the pairwise comparisons of log_2_ ratios between the Δ*rimBK rpsF_10glu_* and Δ*rimBK* translatomes. **(A-C)** Highlighted regions containing significantly (>1.0 log_2_) affected class 1, 2 and 3 genes respectively. **(D)** Genes significantly (>1.0 log_2_) affected by *rpsF* glutamation but not significantly affected by *rimBK* deletion. Highlighted genes are listed in Table S4 and colour-coded according to their COG classifications: yellow = metabolism; green = cellular processes and signalling; blue = information storage and processing; dark grey = poorly characterised.

**Figure S6:**
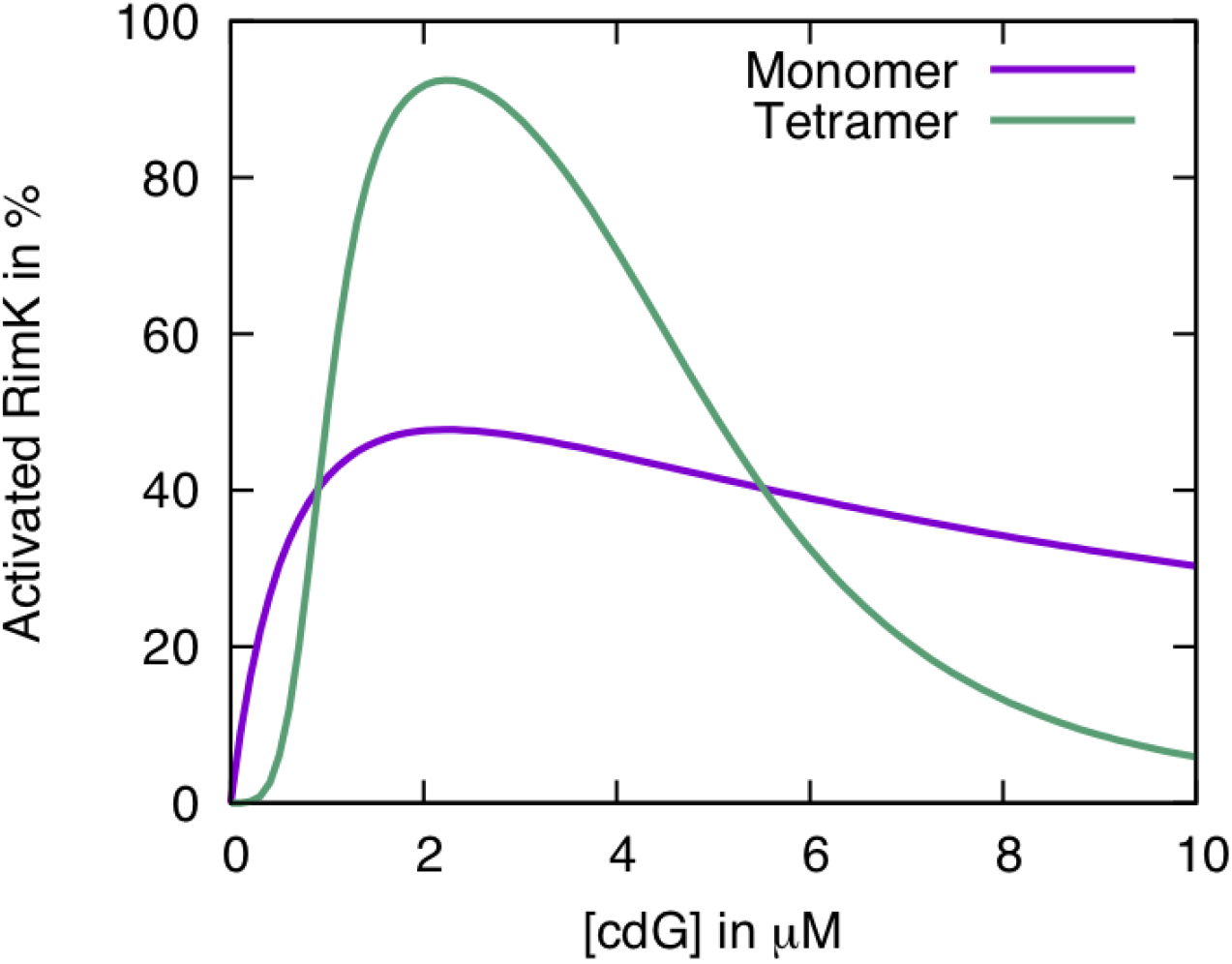
cdG binding to RimA in the presence of RimB could lead to cdG having a dual role in RimK regulation. Our experimental data show that both cdG and RimA increase the ATPase and glutamate activity of RimK in the absence of RimB. A model of RimA and cdG binding to RimK, leading to a state of higher activity, is consistent with this data (Fig S2). In the presence of RimB, however, RimA becomes catalytically active, binding to and modifying cdG. Given the experimental data, a reasonable hypothesis is that this RimA activity promotes, either directly or indirectly, the dissociation from RimK. Here we model the effect of stimulating the RimK.RimA complex by cdG binding to RimK (K_d_ = 1 µM) and its inhibition by binding cdG to RimA (K_d_ = 5 µM) for different assumptions of different complex arrangements for RimK.RimA.

## Supplemental tables

**Table S1.** Kinetic models for RimK ATPase activity.

| Model | Chemical reactions | Constraints |
| --- | --- | --- |
| **Two state**<br>**RimK** | $RimK + RimA \rightleftharpoons RimK^*$<br>$RimK + cdG \rightleftharpoons RimK^*$<br>$RimK + ATP \rightleftharpoons RimK.ATP \rightarrow RimK + ADP + P_i$<br>$RimK^* + ATP \rightleftharpoons RimK^*.ATP \rightarrow RimK^* + ADP + P_i$ | $RimK + RimK^*$<br>$= RimK^{Total}$ |
| **Three state**<br>**RimK** | $RimK + RimA \rightleftharpoons RimK^1$<br>$RimK + cdG \rightleftharpoons RimK^2$<br>$RimK^1 + cdG \rightleftharpoons RimK^2$<br>$RimK^2 + RimA \rightleftharpoons RimK^2$<br>$RimK + ATP \rightleftharpoons RimK.ATP \rightarrow RimK + ADP + P_i$<br>$RimK^1 + ATP \rightleftharpoons RimK^1.ATP \rightarrow RimK^1 + ADP + P_i$<br>$RimK^2 + ATP \rightleftharpoons RimK^2.ATP \rightarrow RimK^2 + ADP + P_i$ | $RimK + RimK^1 + RimK^2$<br>$= RimK^{Total}$<br>$K_{d:RimK.RimA}K_{d:RimK^1.cdG}$<br>$= K_{d:RimK.cdG}K_{d:RimK^2.RimA}$ |
| **Four state**<br>**RimK** | $RimK + RimA \rightleftharpoons RimK^1$<br>$RimK + cdG \rightleftharpoons RimK^2$<br>$RimK^1 + cdG \rightleftharpoons RimK^3$<br>$RimK^2 + RimA \rightleftharpoons RimK^3$<br>$RimK + ATP \rightleftharpoons RimK.ATP \rightarrow RimK + ADP + P_i$<br>$RimK^1 + ATP \rightleftharpoons RimK^1.ATP \rightarrow RimK^1 + ADP + P_i$<br>$RimK^2 + ATP \rightleftharpoons RimK^2.ATP \rightarrow RimK^2 + ADP + P_i$<br>$RimK^3 + ATP \rightleftharpoons RimK^3.ATP \rightarrow RimK^3 + ADP + P_i$ | $RimK + RimK^1 + RimK^2$<br>$+ RimK^3 = RimK^{Total}$<br>$K_{d:RimK.RimA}K_{d:RimK^1.cdG}$<br>$= K_{d:RimK.cdG}K_{d:RimK^2.RimA}$ |

**Table S2 Effect of rimK deletion on relative protein abundance in different media conditions**

Data are shown for all proteins differentially regulated in soluble cell lysates of SBW25 WT compared to Δ*rimK* after 45 minutes exposure to carbon free Rooting Solution at 8 °C, or LB medium at 28 °C. This list represents the proteins shown as coloured spots in Fig 8.

**Table S3 Riboseq data for SBW25 Δ*rimBK***

Genes whose translational activity is greater than log_2_-fold up- or downregulated in SBW25 Δ*rimBK* compared to WT SBW25.

**Table S4 Riboseq data for SBW25 ΔrimBK rpsF_4glu/10glu_**

Genes whose translational activity is greater than log_2_-fold up- or downregulated in SBW25 Δ*rimBK rpsF_4glu/10glu_* compared to Δ*rimBK*.

**Table S5.**
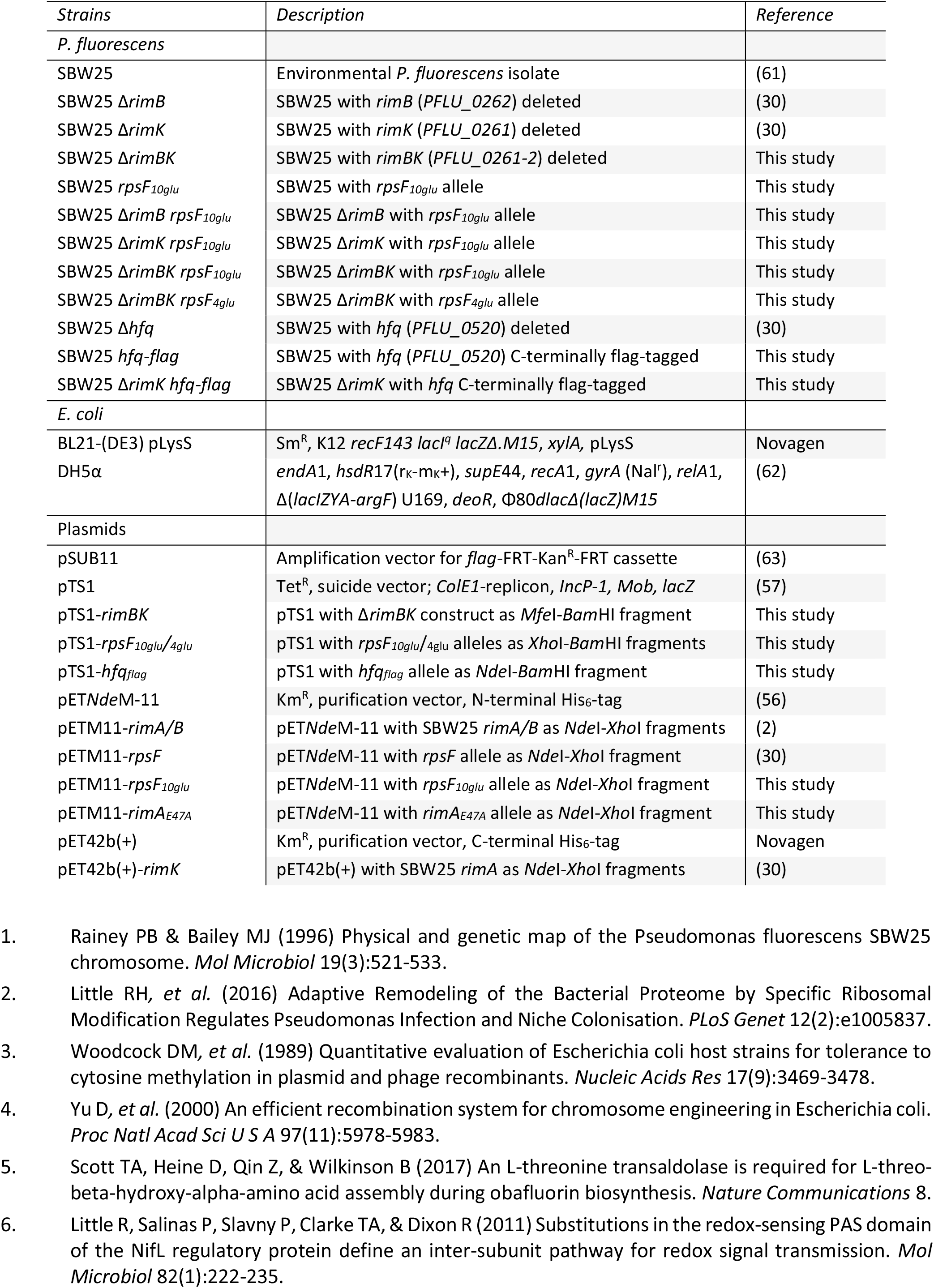
Class II and Class III RimK affected genes

**Table S6.**
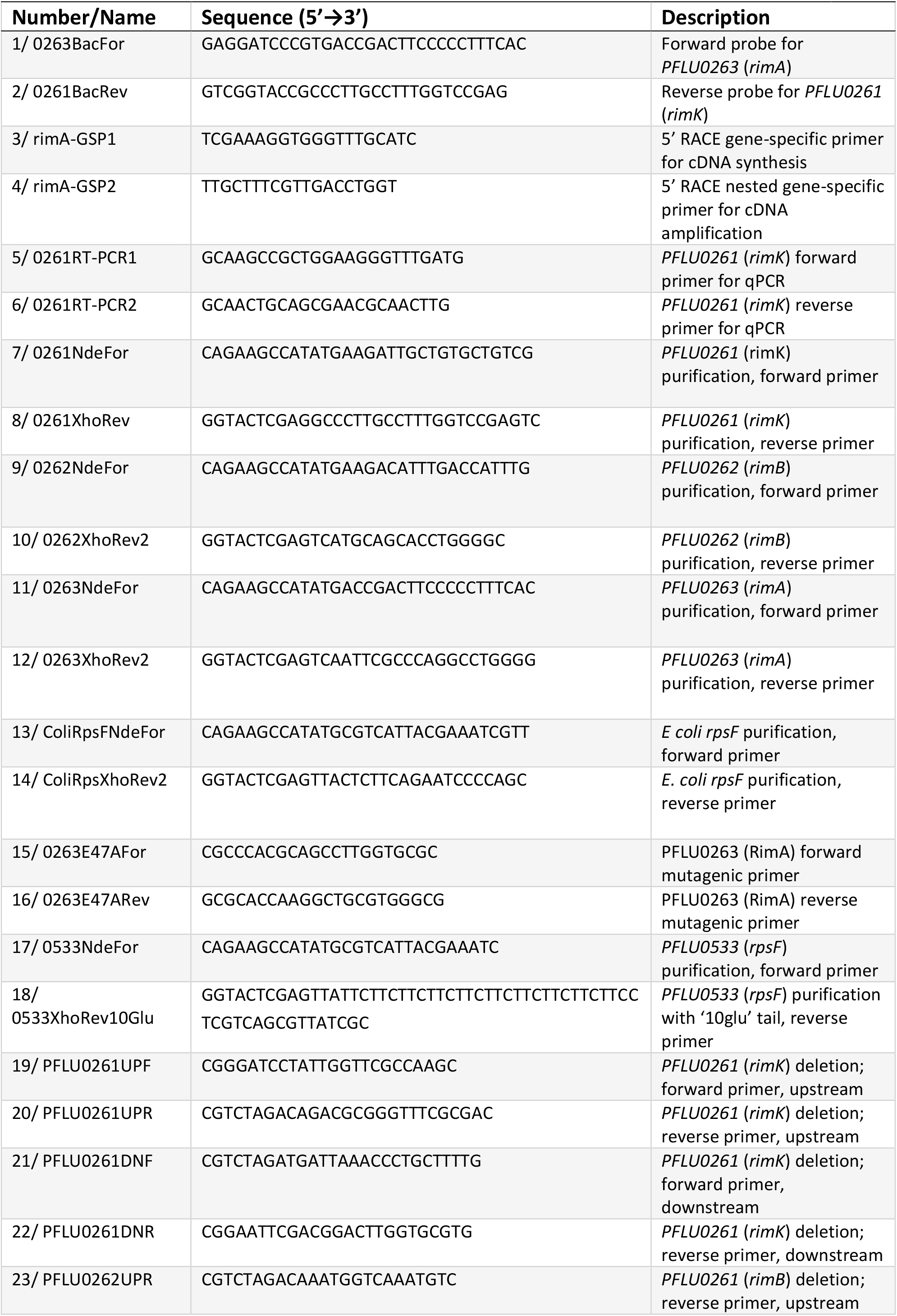
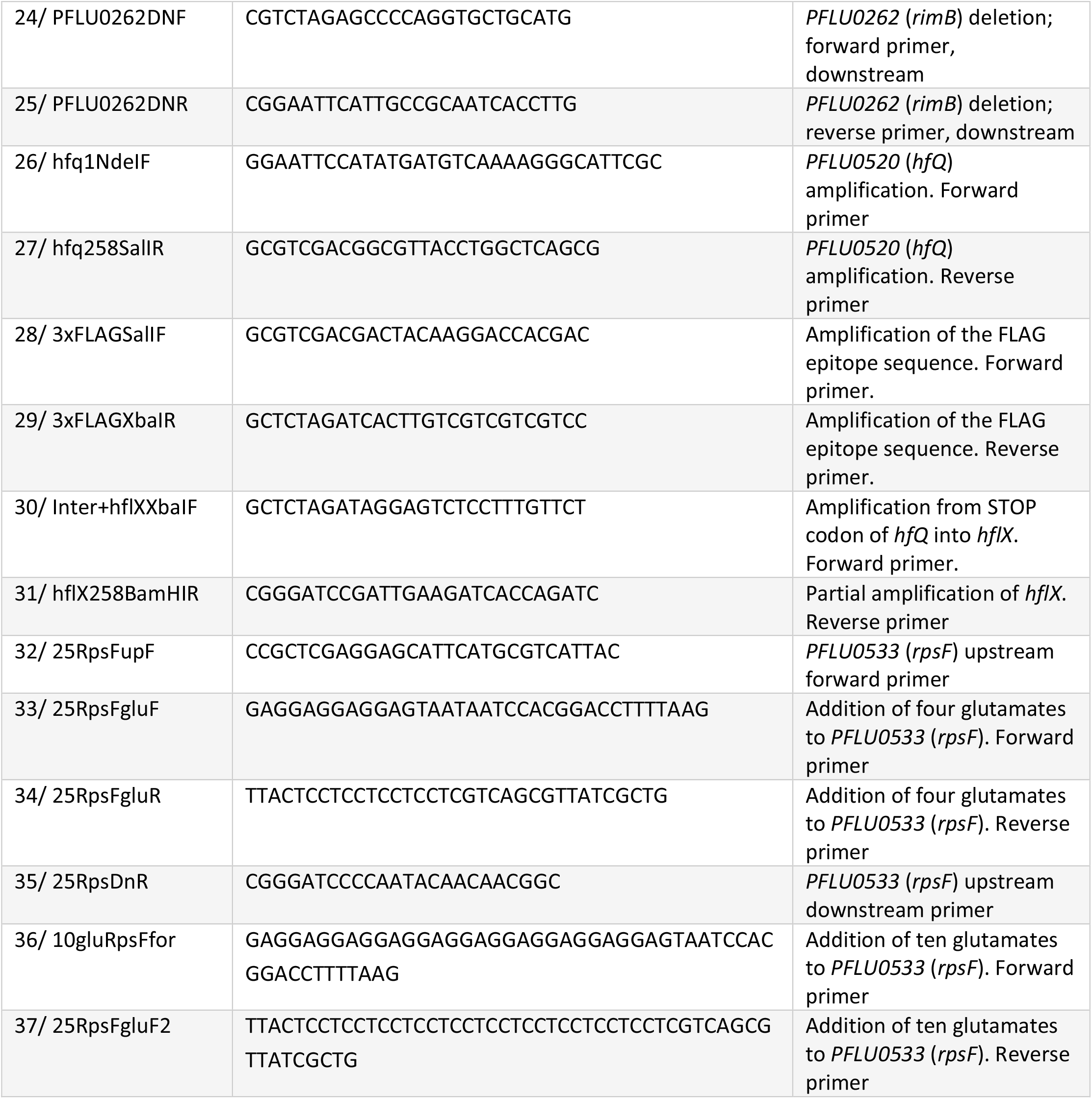
Strains and plasmids used in this study

**Table S7 Primers used in this study**

## References

1. Zhalnina K, et al. (2018) Dynamic root exudate chemistry and microbial substrate preferences drive patterns in rhizosphere microbial community assembly. Nat Microbiol 3(4):470–480.

2. Lundberg DS, et al. (2012) Defining the core Arabidopsis thaliana root microbiome. Nature 488(7409):86–90.

3. Mauchline TH & Malone JG (2017) Life in earth - the root microbiome to the rescue? Curr Opin Microbiol 37:23–28.

4. Alsohim AS, et al. (2014) The biosurfactant viscosin produced by Pseudomonas fluorescens SBW25 aids spreading motility and plant growth promotion. Environ Microbiol 16(7):2267–2281.

5. Chin-A-Woeng TFC, de Priester W, van der Bij AJ, & Lugtenberg BJJ (1997) Description of the Colonization of a Gnotobiotic Tomato Rhizosphere by Pseudomonas fluorescens Biocontrol Strain WCS365, Using Scanning Electron Microscopy. Molecular Plant-Microbe Interactions 10(1):79–86.

6. Lugtenberg BJ, Dekkers L, & Bloemberg GV (2001) Molecular determinants of rhizosphere colonization by Pseudomonas. Annual review of phytopathology 39:461–490.

7. Gal M, Preston GM, Massey RC, Spiers AJ, & Rainey PB (2003) Genes encoding a cellulosic polymer contribute toward the ecological success of Pseudomonas fluorescens SBW25 on plant surfaces. Mol Ecol 12(11):3109–3121.

8. Hinsa SM, Espinosa-Urgel M, Ramos JL, & O’Toole GA (2003) Transition from reversible to irreversible attachment during biofilm formation by Pseudomonas fluorescens WCS365 requires an ABC transporter and a large secreted protein. Mol Microbiol 49(4):905–918.

9. de Weert S, et al. (2006) The two-component colR/S system of Pseudomonas fluorescens WCS365 plays a role in rhizosphere competence through maintaining the structure and function of the outer membrane. FEMS microbiology ecology 58(2):205–213.

10. Ryu CM, et al. (2003) Bacterial volatiles promote growth in Arabidopsis. Proc Natl Acad Sci U S A 100(8):4927–4932.

11. Loper JE, et al. (2012) Comparative genomics of plant-associated Pseudomonas spp.: insights into diversity and inheritance of traits involved in multitrophic interactions. PLoS Genet 8(7):e1002784.

12. Klee HJ, Hayford MB, Kretzmer KA, Barry GF, & Kishore GM (1991) Control of ethylene synthesis by expression of a bacterial enzyme in transgenic tomato plants. The Plant cell 3(11):1187–1193.

13. Dimkpa CO, Merten D, Svatos A, Buchel G, & Kothe E (2009) Metal-induced oxidative stress impacting plant growth in contaminated soil is alleviated by microbial siderophores. Soil Biol Biochem 41(1):154–162.

14. Haas D & Defago G (2005) Biological control of soil-borne pathogens by fluorescent pseudomonads. Nat Rev Microbiol 3(4):307–319.

15. Compant S, Clement C, & Sessitsch A (2010) Plant growth-promoting bacteria in the rhizo- and endosphere of plants: Their role, colonization, mechanisms involved and prospects for utilization. Soil Biology and Biochemistry 42:669–678.

16. Ramachandran VK, East AK, Karunakaran R, Downie A, & Poole PS (2011) Adaptation of Rhizobium leguminosarum to pea, alfalfa and sugar beet rhizospheres investigated by comparative transcriptomics. Genome Biol 12(10):R106.

17. Campilongo R, et al. (2017) One ligand, two regulators and three binding sites: How KDPG controls primary carbon metabolism in Pseudomonas. PLoS Genet 13(6):e1006839.

18. Grenga L, et al. (2017) Analysing the complex regulatory landscape of Hfq - an integrative, multi-omics approach. Front Microbiol (8):1784.

19. Martinez-Granero F, et al. (2014) Identification of flgZ as a flagellar gene encoding a PilZ domain protein that regulates swimming motility and biofilm formation in Pseudomonas. PLoS One 9(2):e87608.

20. Jenal U, Reinders A, & Lori C (2017) Cyclic di-GMP: second messenger extraordinaire. Nat Rev Microbiol 15(5):271–284.

21. Barahona E, et al. (2011) Pseudomonas fluorescens F113 Mutant with Enhanced Competitive Colonization Ability and Improved Biocontrol Activity against Fungal Root Pathogens. Appl Environ Microbiol 77(15):5412–5419.

22. Little RH, et al. (2019) Differential Regulation of Genes for Cyclic-di-GMP Metabolism Orchestrates Adaptive Changes During Rhizosphere Colonization by Pseudomonas fluorescens. Frontiers in Microbiology 10, 1089.

23. O’Neal L, Akhter S, & Alexandre G (2019) A PilZ-Containing Chemotaxis Receptor Mediates Oxygen and Wheat Root Sensing in Azospirillum brasilense. Front Microbiol 10:312.

24. Ramos-Gonzalez MI, et al. (2016) Genetic Dissection of the Regulatory Network Associated with High c-di-GMP Levels in Pseudomonas putida KT2440. Front Microbiol 7:1093.

25. Grenga L, Little RH, & Malone JG (2017) Quick Change - post-transcriptional regulation in Pseudomonas. FEMS Microbiol Lett. 364 (14).

26. Valentini M & Filloux A (2016) Biofilms and Cyclic di-GMP (c-di-GMP) Signaling: Lessons from Pseudomonas aeruginosa and Other Bacteria. J Biol Chem 291(24):12547–12555.

27. Silby MW, et al. (2009) Genomic and genetic analyses of diversity and plant interactions of Pseudomonas fluorescens. Genome Biol 10(5):R51.

28. Moscoso JA, et al. (2014) The diguanylate cyclase SadC is a central player in Gac/Rsm-mediated biofilm formation in Pseudomonas aeruginosa. J Bacteriol 196(23):4081–4088.

29. Moscoso JA, Mikkelsen H, Heeb S, Williams P, & Filloux A (2011) The Pseudomonas aeruginosa sensor RetS switches type III and type VI secretion via c-di-GMP signalling. Environ Microbiol 13(12):3128–3138.

30. Little RH, et al. (2016) Adaptive Remodeling of the Bacterial Proteome by Specific Ribosomal Modification Regulates Pseudomonas Infection and Niche Colonisation. PLoS Genet 12(2):e1005837.

31. Moller T, et al. (2002) Hfq: a bacterial Sm-like protein that mediates RNA-RNA interaction. Mol Cell 9(1):23–30.

32. Maki K, Uno K, Morita T, & Aiba H (2008) RNA, but not protein partners, is directly responsible for translational silencing by a bacterial Hfq-binding small RNA. Proc Natl Acad Sci U S A 105(30):10332–10337.

33. Jorgensen MG, et al. (2012) Small regulatory RNAs control the multi-cellular adhesive lifestyle of Escherichia coli. Mol Microbiol 84(1):36–50.

34. Sonnleitner E BU (2014) Regulation of Hfq by the RNA CrcZ in Pseudomonas aeruginosa carbon catabolite repression. Plos Genetics 10(6):e1004440.

35. Sobrero P, et al. (2012) Quantitative proteomic analysis of the Hfq-regulon in Sinorhizobium meliloti 2011. PLoS One 7(10):e48494.

36. Gao M, Barnett MJ, Long SR, & Teplitski M (2010) Role of the Sinorhizobium meliloti global regulator Hfq in gene regulation and symbiosis. Molecular plant-microbe interactions: MPMI 23(4):355–365.

37. Sonnleitner E, et al. (2003) Reduced virulence of a hfq mutant of Pseudomonas aeruginosa O1. Microb Pathog 35(5):217–228.

38. Irie Y, et al. (2010) Pseudomonas aeruginosa biofilm matrix polysaccharide Psl is regulated transcriptionally by RpoS and post-transcriptionally by RsmA. Mol Microbiol 78(1):158–172.

39. Brencic A & Lory S (2009) Determination of the regulon and identification of novel mRNA targets of Pseudomonas aeruginosa RsmA. Mol Microbiol 72(3):612–632.

40. Kino K, Arai T, & Arimura Y (2011) Poly-alpha-glutamic acid synthesis using a novel catalytic activity of RimK from Escherichia coli K-12. Appl Environ Microbiol 77(6):2019–2025.

41. Kaczanowska M & Ryden-Aulin M (2007) Ribosome biogenesis and the translation process in Escherichia coli. Microbiol Mol Biol Rev 71(3):477–494.

42. Shen J, Meldrum A, & Poole K (2002) FpvA receptor involvement in pyoverdine biosynthesis in Pseudomonas aeruginosa. J Bacteriol 184(12):3268–3275.

43. Nait Chabane Y, et al. (2014) Characterisation of pellicles formed by Acinetobacter baumannii at the air-liquid interface. PLoS One 9(10):e111660.

44. Poole P, Ramachandran V, & Terpolilli J (2018) Rhizobia: from saprophytes to endosymbionts. Nat Rev Microbiol 16(5):291–303.

45. Schulz S, et al. (2015) Elucidation of sigma factor-associated networks in Pseudomonas aeruginosa reveals a modular architecture with limited and function-specific crosstalk. PLoS Pathog 11(3):e1004744.

46. Lindenberg S, Klauck G, Pesavento C, Klauck E, & Hengge R (2013) The EAL domain protein YciR acts as a trigger enzyme in a c-di-GMP signalling cascade in E. coli biofilm control. The EMBO journal 32(14):2001–2014.

47. Loveland AB & Korostelev AA (2018) Structural dynamics of protein S1 on the 70S ribosome visualized by ensemble cryo-EM. Methods 137:55–66.

48. Duval M, et al. (2013) Escherichia coli ribosomal protein S1 unfolds structured mRNAs onto the ribosome for active translation initiation. PLoS Biol 11(12):e1001731.

49. Sukhodolets MV & Garges S (2003) Interaction of Escherichia coli RNA polymerase with the ribosomal protein S1 and the Sm-like ATPase Hfq. Biochemistry 42(26):8022–8034.

50. Kambara TK, Ramsey KM, & Dove SL (2018) Pervasive Targeting of Nascent Transcripts by Hfq. Cell Rep 23(5):1543–1552.

51. Carmichael GG, Weber K, Niveleau A, & Wahba AJ (1975) The host factor required for RNA phage Qbeta RNA replication in vitro. Intracellular location, quantitation, and purification by polyadenylate-cellulose chromatography. J Biol Chem 250(10):3607–3612.

52. Wahba AJ, et al. (1974) Subunit I of G beta replicase and 30 S ribosomal protein S1 of Escherichia coli. Evidence for the identity of the two proteins. J Biol Chem 249(10):3314–3316.

53. Inouye H, Pollack Y, & Petre J (1974) Physical and functional homology between ribosomal protein S1 and interference factor i. Eur J Biochem 45(1):109–117.

54. Kang WK, Icho T, Isono S, Kitakawa M, & Isono K (1989) Characterization of the gene rimK responsible for the addition of glutamic acid residues to the C-terminus of ribosomal protein S6 in Escherichia coli K12. Molecular & general genetics: MGG 217(2-3):281–288.

55. Miller JH (1972) Experiments in molecular genetics. Cold Spring Harbor Laboratory, Cold Spring Harbor, New York:352–355.

56. Little R, Salinas P, Slavny P, Clarke TA, & Dixon R (2011) Substitutions in the redox-sensing PAS domain of the NifL regulatory protein define an inter-subunit pathway for redox signal transmission. Mol Microbiol 82(1):222–235.

57. Scott TA, Heine D, Qin Z, & Wilkinson B (2017) An L-threonine transaldolase is required for L-threo-beta-hydroxy-alpha-amino acid assembly during obafluorin biosynthesis. Nature Communications 8.

58. King EO, Ward MK, & Raney DE (1954) Two simple media for the demonstration of pyocyanin and fluorescin. The Journal of laboratory and clinical medicine 44(2):301–307.

59. Oh E BA, Sandikci A, Huber D, Chaba R, Gloge F, Nichols RJ, Typas A, Gross CA, Kramer G, Weissman JS, Bukau B. (2011) Selective ribosome profiling reveals the cotranslational chaperone action of trigger factor in vivo. Cell 147(6):1295–1308.

60. Becker AH, Oh E, Weissman JS, Kramer G, & Bukau B (2013) Selective ribosome profiling as a tool for studying the interaction of chaperones and targeting factors with nascent polypeptide chains and ribosomes. Nature protocols 8(11):2212–2239.

61. Rainey PB & Bailey MJ (1996) Physical and genetic map of the Pseudomonas fluorescens SBW25 chromosome. Mol Microbiol 19(3):521–533.

62. Woodcock DM, et al. (1989) Quantitative evaluation of Escherichia coli host strains for tolerance to cytosine methylation in plasmid and phage recombinants. Nucleic Acids Res 17(9):3469–3478.

63. Yu D, et al. (2000) An efficient recombination system for chromosome engineering in Escherichia coli. Proc Natl Acad Sci U S A 97(11):5978–5983.

